# Iron oxidation by a fused cytochrome-porin common to diverse iron-oxidizing bacteria

**DOI:** 10.1101/228056

**Authors:** Jessica L. Keffer, Sean M. McAllister, Arkadiy Garber, Beverly J. Hallahan, Molly C. Sutherland, Sharon Rozovsky, Clara S. Chan

## Abstract

Iron (Fe) oxidation is one of Earth’s major biogeochemical processes, key to weathering, soil formation, water quality, and corrosion. However, our understanding of microbial contribution is limited by incomplete knowledge of microbial iron oxidation mechanisms, particularly in neutrophilic iron-oxidizers. The genomes of many, diverse iron-oxidizers encode a homolog to an outer-membrane cytochrome (Cyc2) shown to oxidize iron in two acidophiles. Phylogenetic analyses show Cyc2 sequences from neutrophiles cluster together, suggesting a common function, though this function has not been verified in these organisms. Therefore, we investigated the iron oxidase function of heterologously expressed Cyc2 from a neutrophilic iron-oxidizer *Mariprofundus ferrooxydans* PV-1. Cyc2PV-1 is capable of oxidizing iron, and its redox potential is 208 ± 20 mV, consistent with the ability to accept electrons from Fe^2+^ at neutral pH. These results support the hypothesis that Cyc2 functions as an iron oxidase in neutrophilic iron-oxidizing organisms. Sequence analysis and modeling reveal the entire Cyc2 family share a unique fused cytochrome-porin structure, with a defining consensus motif in the cytochrome region. Based on structural analyses, we predict that the monoheme cytochrome Cyc2 specifically oxidizes dissolved Fe^2+^, in contrast to multiheme iron oxidases, which may oxidize solid Fe(II). With our results, there is now functional validation for diverse representatives of Cyc2 sequences. We present a comprehensive Cyc2 phylogenetic tree and offer a roadmap for identifying *cyc2/*Cyc2 homologs and interpreting their function. The occurrence of *cyc2* in many genomes beyond known iron-oxidizers presents the possibility that microbial iron oxidation may be a widespread metabolism.

**Importance:** Iron is practically ubiquitous across Earth’s environments, central to both life and geochemical processes, which depend heavily on the redox state of iron. Although iron oxidation, or “rusting,” can occur abiotically at near neutral pH, we find neutrophilic iron-oxidizing bacteria (FeOB) are widespread, including in aquifers, sediments, hydrothermal vents, pipes, and water treatment systems. FeOB produce highly reactive Fe(III) oxyhydroxides that bind a variety of nutrients and toxins, thus these microbes are likely a controlling force in iron and other biogeochemical cycles. There has been mounting evidence that Cyc2 functions as an iron oxidase in neutrophiles, but definitive proof of its function has long eluded us. This work provides conclusive biochemical evidence of iron oxidation by Cyc2 from neutrophiles. Cyc2 is common to a wide variety of iron-oxidizers, including acidophilic and phototrophic iron-oxidizers, suggesting that this fused cytochrome-porin structure is especially well-adapted for iron oxidation.

## Introduction

Iron (Fe) oxidation occurs in virtually all near-surface environments, producing highly reactive iron oxyhydroxides that often control the fate of carbon, phosphorous, and other metals (1). It is often assumed that abiotic reactions are the primary mechanisms of iron oxidation, particularly at near-neutral pH. However, iron-oxidizing microbes are increasingly observed in a wide range of environments, especially dark, neutral pH environments such as aquifers, soils, sediments, hydrothermal vents, and water treatment systems (2–7). Neutrophilic iron-oxidizers thrive at anoxic-oxic interfaces where they can outcompete abiotic iron oxidation rates at low oxygen concentrations (8). Iron-oxidizing microbes can grow by coupling iron oxidation to the reduction of oxygen or nitrate, using this energy to fuel carbon fixation and biomass production (2, 9, 10), but in many iron-replete environments, we lack a clear understanding of the extent of microbial iron oxidation. To address this, we need to confidently identify iron oxidation mechanisms. Yet, unlike other major microbial metabolisms, we have relatively incomplete knowledge of iron oxidation pathways (11–13).

Rising interest in iron-oxidizing microbes has resulted in a surge of iron-oxidizer sequencing, including isolate genomes, single cell genomes, metagenomes, and metatranscriptomes (5, 7, 9, 14–20), enabling us to search for the genes involved in microbial iron oxidation using genome mining. All sequenced genomes of the known neutrophilic chemolithoautotrophic iron-oxidizing bacteria (FeOB), the marine Zetaproteobacteria (*Mariprofundus* spp., *Ghiorsea* spp.) and freshwater Gallionellaceae (*Gallionella* spp., *Sideroxydans* spp., and *Ferriphaselus* spp.), have homologs to *cyc2* (7, 21–25), which encodes an iron-oxidizing outer membrane cytochrome first characterized in *Acidithiobacillus ferrooxidans* (26, 27). Many of these FeOB are obligate iron-oxidizers that lack other apparent iron oxidase candidates. A second potential iron oxidase gene, *mtoA,* is found in a few Gallionellaceae and *Thiomonas* genomes (21, 28, 29), and functional and genetic information supports the role of MtoA and its homolog PioA in iron oxidation (30–32). However, relatively few FeOB genomes contain *mtoA,* and *pioA* is limited to phototrophic organisms, suggesting that Cyc2 is potentially a more widespread iron oxidase.

Recently, McAllister *et al*. presented the phylogeny of 634 unique Cyc2 homologs (7), which resolved into three distinct clusters. Two of the clusters each contain a Cyc2 homolog with verified iron-oxidizing activity - *Acidithiobacillus ferrooxidans* Cyc2 (27) in Cluster 2 and *Leptospirillum* sp. Cyt_572_ (33) in Cluster 3. Both of these organisms are acidophilic iron-oxidizers. Cluster 1 consists largely of the well-known neutrophilic iron-oxidizers, including the Zetaproteobacteria, Gallionellaceae, and iron-oxidizing *Chlorobium* spp. This cluster is well-supported and these sequences are among the closest homologs to one another despite the taxonomic distance between these organisms (7). A common iron oxidation pathway for both neutrophiles and acidophiles might not be expected, due to the drastically different redox potential of Fe(II)/Fe(III) at acidic versus neutral pH (770 mV at pH 2 vs. 0 ± 100 mV at pH 7 (11, 34–36)), but a conserved protein structure could suggest a shared function. To be more confident that Cyc2 is an iron oxidase in a wide range of iron-oxidizers, we need biochemical verification of Cyc2 activity from neutrophilic chemolithotrophic FeOB.

Our main objective was to demonstrate iron-oxidizing function for Cyc2 from the well-supported Cluster 1 containing neutrophilic FeOB. We chose the Cluster 1 Cyc2 from *Mariprofundus ferrooxydans* PV-1, an obligate iron-oxidizer, with *cyc2* as the only identified iron oxidase gene candidate in its genome (7, 37–39). Environmental metatranscriptomics of marine iron-oxidizing microbial mats dominated by Zetaproteobacteria, including *Mariprofundus* spp. found high expression of *cyc2*, along with evidence that *cyc2* expression may be regulated in response to Fe(II), suggesting utility for *cyc2* in an iron-oxidizing environment (7, 10). Proteomics of *M. ferrooxydans* PV-1 showed that Cyc2 was expressed, and a membrane complex containing Cyc2 possessed ferrocyanide oxidation activity (39). While the *A. ferrooxidans* Cyc2 characterization was performed in the native organism (27), neutrophilic FeOB are challenging to grow in quantities sufficient for protein assays, so we took a heterologous expression approach. We first performed structure-function predictions to inform the design of the expression constructs, which we then expressed in *E. coli*. To test the iron oxidase function, we expressed Cyc2PV-1 under native conditions and performed iron oxidation assays and redox potential measurements of Cyc2PV-1 enriched by purification chromatography.

We integrate structural, functional, and phylogenetic insights to explore the function of a wider range of Cyc2 homologs, including strategies for correctly identifying Cyc2 homologs and interpreting Cyc2 iron oxidizing function.

## Results & Discussion

### Cyc2 is a predicted outer membrane cytochrome fused to a porin

We first performed a primary and secondary structure analysis to better understand the functional parts of Cyc2PV-1 prior to expression. Cyc2 starts with a signal peptide, predicted by SignalP (40) and has a single - CXXCH-heme-binding motif (41), placing this cytochrome in the c-type family. Fused to the cytochrome domain is a beta-barrel porin, based on the presence of 16 beta strands predicted by PSIPRED (42, 43) (Fig. S1) and homology matching by HHpred (44, 45). The porin-encoding segment has low sequence homology but high structural homology to the outer membrane phosphate-selective porins OprO and OprP (PDB structures 4RJW and 2O4V (46, 47)), based on structural homology predictions by I-TASSER, MODELLER, and Phyre2 (48–51) (Fig. 1). We performed HHpred structural predictions for a diverse set of Cyc2 sequences from McAllister *et al*. (7), and all analyzed sequences matched to the same tertiary structure. Thus, all Cyc2 sequences are predicted to have the same structure: a 16-stranded outer membrane porin with an N-terminal cytochrome domain (Fig. 1).

**Figure 1.**
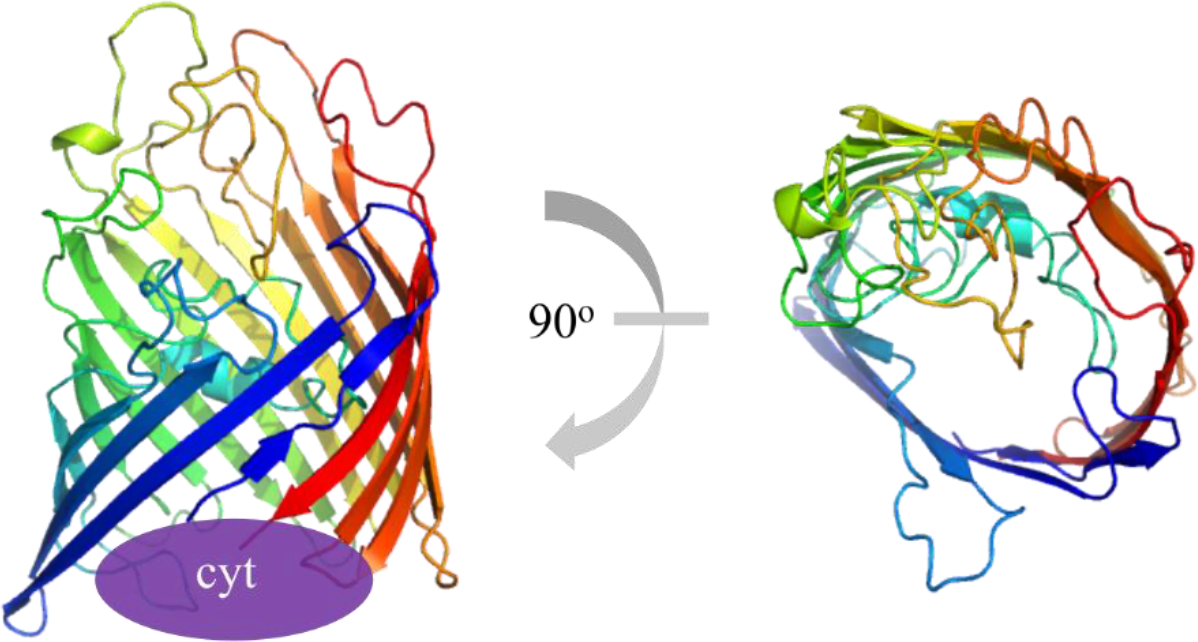
I-TASSER (49) model of Cyc2PV-1. Cyc2 is predicted to be a 16-stranded porin with a fused N-terminal cytochrome domain. The Cyc2 cytochrome (purple oval) is connected to the N-terminal (blue strand) end of the porin. The view on the right is rotated 90° towards the viewer to show the internal pore of the porin but does not contain the cytochrome.

The combination of signal peptide and beta-barrel structure suggests Cyc2 is localized to the outer membrane, as expected for a porin (52). This location is consistent with previous observations that iron oxidation occurs at the cell surface, preventing internal iron oxyhydroxide encrustation (53, 54). Porins typically possess short periplasmic turns and longer extracellular loops, and have both the N- and C-termini in the periplasmic space (55). This standard orientation applied to Cyc2 would suggest that the cytochrome domain of Cyc2 resides on the periplasmic side of the barrel, likely as a plug at the opening of the beta barrel pore.

### Strategy for expression of fused cytochrome-porin

Over-expression strategies for porin proteins typically include directing expression to the cytoplasm where they form inclusion bodies that can be purified, and the porin is subsequently refolded (56). However, c-type cytochromes must be matured in the oxidizing environment of the periplasm to ensure covalent attachment of the heme (57). Thus, to ensure proper folding and localization to the outer membrane, our expression strategies were restricted to native conditions, at the expense of yield.

To maintain these functional parts during expression in *E. coli* C41, we synthesized a codon-optimized gene construct, and replaced the PV-1 signal peptide with the signal peptide from *E. coli* OmpA (56). To assist with detection, we placed a *Strep*-tag II at the C-terminus where it would likely not affect cytochrome maturation (Fig. S2A). For purification, we additionally placed a linker, TEV protease cleavage site, and octa-histidine tag (His-tag) following the *Strep*-tag II (Fig. 2A). In addition, we tested independent expression of the cytochrome domain and the porin domain, and while the porin was expressed, the cytochrome was not (data not shown), suggesting contacts within the full structure play a role in its stability.

**Figure 2.**
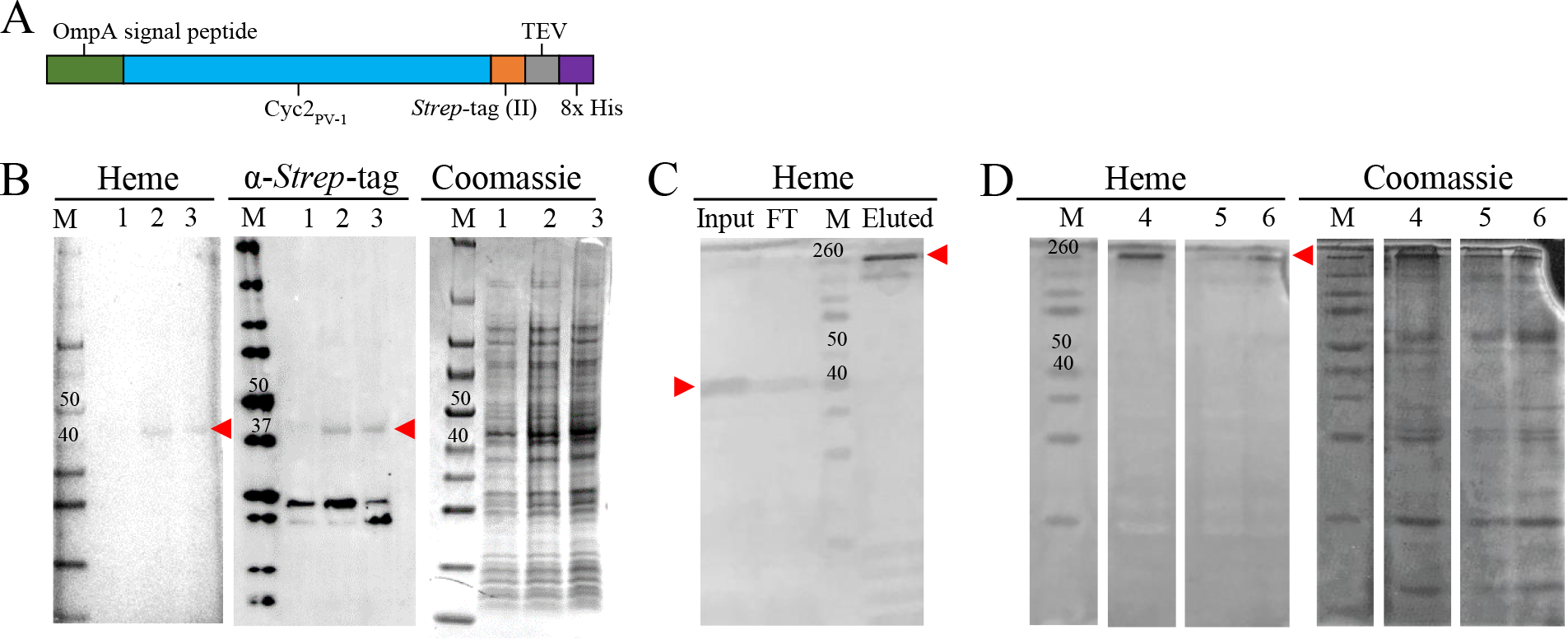
Constructs and expression of Cyc2PV-1. Cyc2PV-1 is marked with a red arrowhead, protein ladder is labeled with an M (Spectra Broad Range on heme and Coomassie, WesternC on α-*Strep*-tag Western blots), and relevant band sizes are labeled in kDa. (A) Schematic of gene construct for expression in *E. coli.* (B) Representative stained SDS-PAGE gels showing Cyc2PV-1 expression in *E. coli*: 1) uninduced, 2) induced, 3) lysed and induced. Smaller bands visible on the *Strep*-tag Western blots are non-specific. (C) Heme-stained gel of fractions during His-tag purification. After elution, Cyc2PV-1 migrates in a high-molecular weight complex. (D) Stained SDS-PAGE gels showing Cyc2PV-1 enrichment fraction: 4) after His-tag purification, 5) after cleavage of His-tag, and 6) after concentration for assays. Uncropped gels shown in Figure S2C.

### *Cyc2PV-1 was produced by heterologous expression in* Escherichia coli

We confirmed production of Cyc2PV-1 in total *E. coli* lysate by immunoblotting with antibodies specific to the C-terminal *Strep*-tag II, as Cyc2_PV-1_ could not be identified solely by Coomassie staining (construct with both *Strep*- and His-tag Fig. 2B; construct with *Strep*-tag only Fig. S2A). We confirmed this same protein contained heme. We were unsuccessful at purification using only the *Strep*-tag, but the His-tag could successfully be used for enrichment of Cyc2_PV-1_ from *E. coli* extracts. Prior to enrichment, Cyc2_PV-1_ ran at its apparent molecular weight and was the only heme-containing protein present. After enrichment, Cyc2_PV-1_ no longer ran true to size, and instead appeared as a high molecular weight band, possibly due to the purification conditions (high salt, high imidazole, and increased protein concentration) (Fig. 2CD; Fig. S2B).

We confirmed Cyc2_PV-1_ is an outer membrane protein by using a sucrose gradient and ultracentrifugation to separate membrane components (58, 59). Cyc2_PV-1_ was found in the outer membrane fraction, in agreement with the predictions from the bioinformatic analysis (Fig. 3A). We isolated the band corresponding to Cyc2_PV-1_ from the outer membrane fraction on a heme-stained SDS-PAGE gel. Tandem mass spectrometry analysis of this band confirmed its identity as Cyc2_PV-1_, as nearly 60% of the protein sequence was detected, including regions of the cytochrome domain, the porin domain, and the *Strep*-tag II (Fig. 3B; Supplementary Table 1).

**Figure 3.**
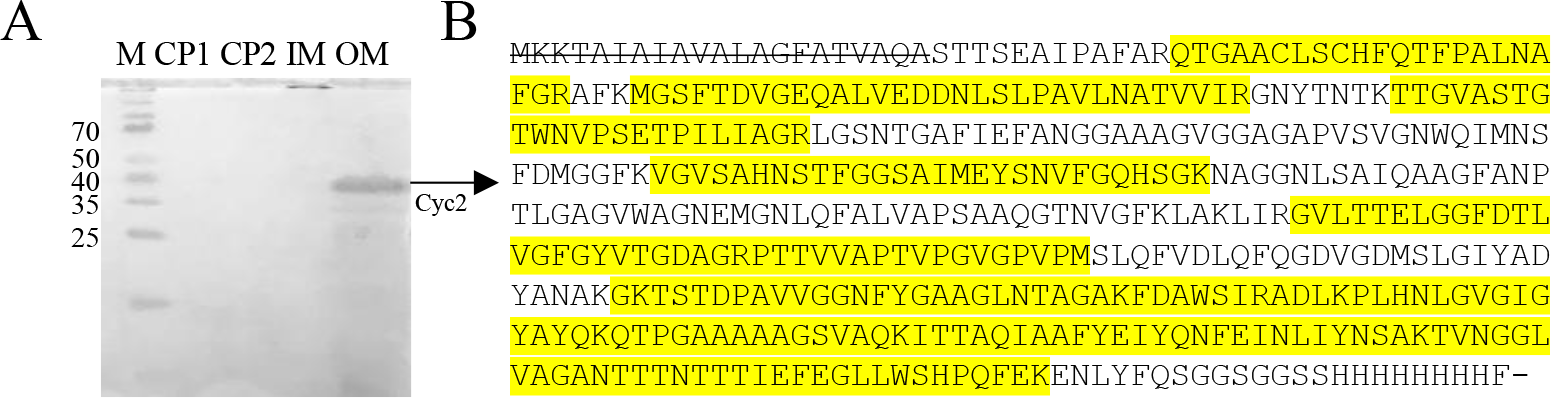
Cyc2_PV-1_ is located in the outer membrane. (A) Heme-stained SDS-PAGE gel of fractionated *E. coli* (CP1, CP2 - cytoplasmic proteins, IM - inner membranes, OM - outer membranes, M - Spectra Broad Range protein ladder with relevant bands in kDa). Cyc2_PV-1_ was found only in the outer membrane fraction (OM), and no other heme-containing proteins were seen in the OM. Coomassie-stained gel in Figure S3. (B) Highlighted yellow peptides observed in tandem MS/MS confirmed the identity of the OM heme-stained band as Cyc2_PV-1_, with nearly 60% of the protein detected. The signal peptide is crossed out, as it is not present in the mature protein.

### Cyc2_PV-1_ has a distinct heme spectrum

We isolated inner and outer membranes from both *E. coli* expressing full-length Cyc2_PV-1_ and *E. coli* expressing only the porin domain of Cyc2_PV-1_. The ultraviolet-visible (UV-Vis) absorbance spectra of these two *E. coli* membrane fractions were compared to identify the heme signal from Cyc2_PV-1_ (Fig. 4). The inner membranes of *E. coli* expressing either Cyc2_PV-1_ or porin-domain only were virtually identical when analyzed by UV-Vis spectroscopy, showing a heme Soret peak at 415 nm (Fig. 4, dashed black and orange spectra). These peaks represent *E. coli’s* native heme-containing proteins. In contrast, there are no native outer membrane heme proteins in *E. coli* (60). Indeed, only the outer membranes from *E. coli* expressing full-length Cyc2_PV-1_ had heme (Fig. 4, solid orange spectra), with a Soret peak at 410 nm. Thus, the Soret peak at 410 nm is indicative of Cyc2_PV-1_ in our system and can be used to spectroscopically detect Cyc2_PV-1_.

**Figure 4.**
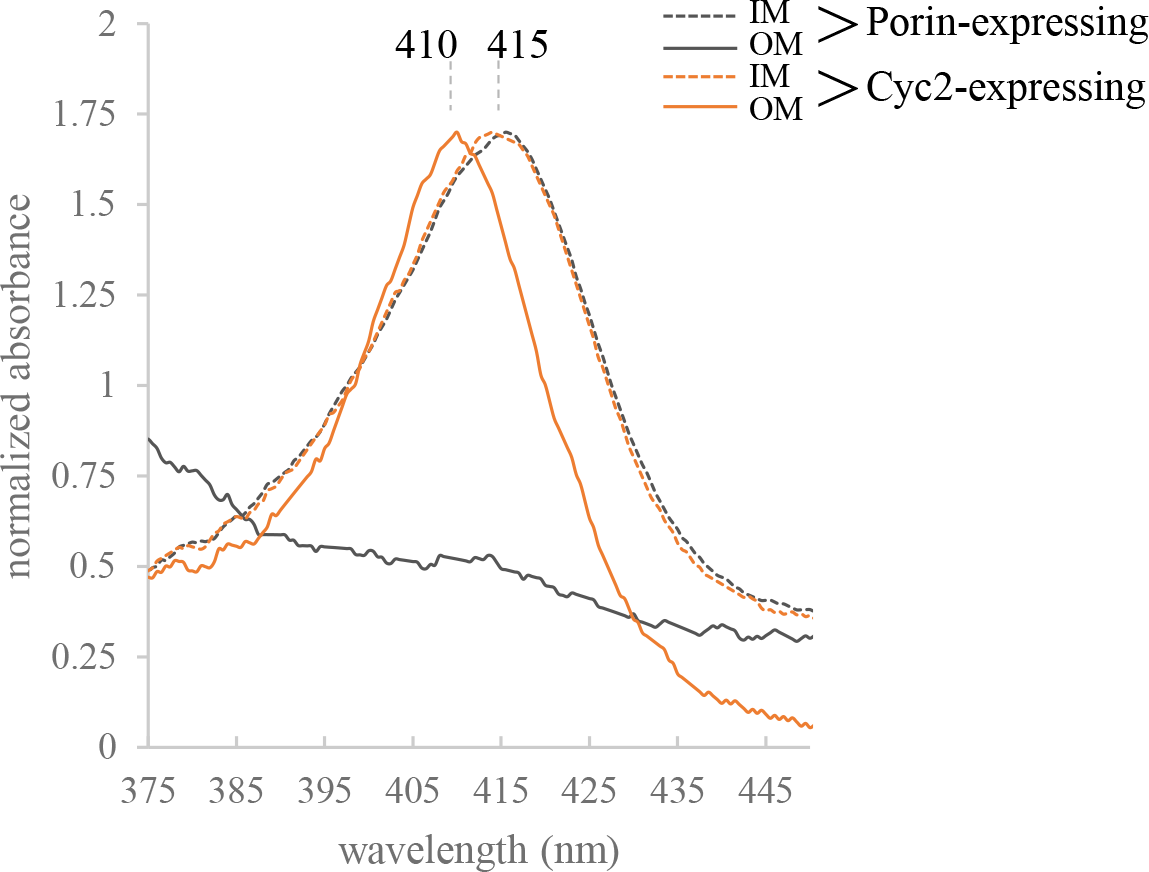
Cyc2_PV-1_ has a distinctive heme Soret peak of 410 nm. UV-Vis spectra of inner (IM) and outer membranes (OM) obtained from a sucrose gradient of *E. coli* expressing either full-length Cyc2_PV-1_ or only the porin-domain of Cyc2_PV-1_. Cyc2_PV-1_ OM had heme (orange), while Porin OM did not (black), and the heme signal was distinct from other heme proteins in *E. coli* (IM, dashed black and orange). Heme-stained and Coomassie-stained SDS-PAGE gels and *Strep*-tag Western blot shown in Figure S3.

### Cyc2_PV-1_ is an iron oxidase

We assayed the iron oxidation capacity of enriched Cyc2_PV-1_ by UV-Vis spectroscopy under anaerobic conditions after removal of the His-tag by TEV protease cleavage. While Cyc2_PV-1_ is not pure, it is the only heme-containing protein detected, and the UV-Vis methods utilized here rely only on the heme spectra. Importantly, the heme spectrum of enriched Cyc2_PV-1_ is identical to the heme spectrum of outer membranes from *E. coli* expressing Cyc2_PV-1_, where Cyc2_PV-1_ was established as the only heme-containing protein (Fig. 4). Cyc2_PV-1_ could be reduced with sodium dithionite (Fig. 5A), demonstrated by the shift of the Soret peak from 410 nm to 427 nm and the appearance of α and β peaks at 560 nm and 530 nm, respectively. Addition of an oxidizing agent to enriched Cyc2_PV-1_ caused no changes, indicating that Cyc2_PV-1_ was in the oxidized state as purified (Fig. 5A). These assays demonstrated Cyc2_PV-1_ was redox active and so we tested its capacity to oxidize Fe(II).

**Figure 5.**
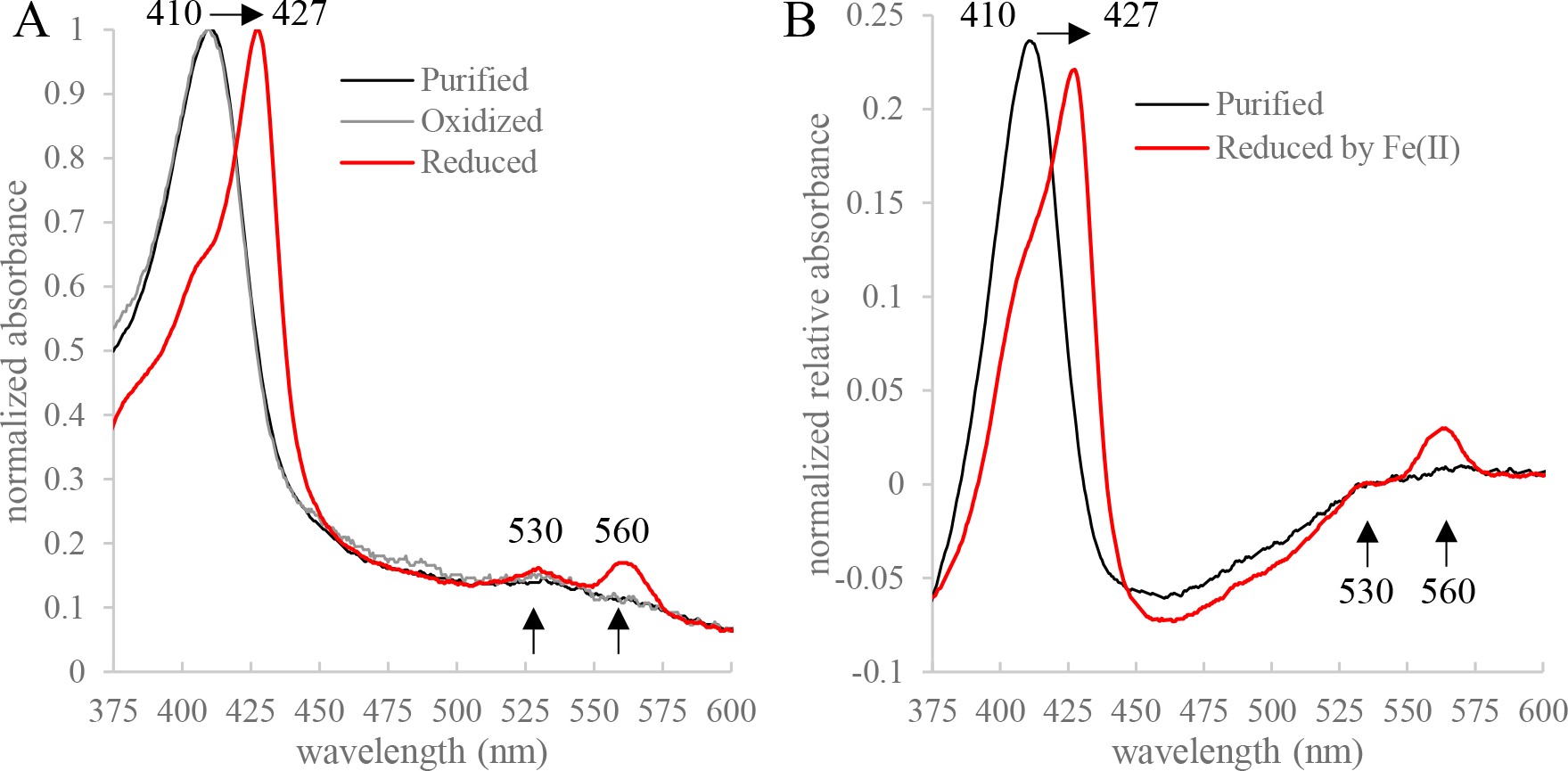
Cyc2_PV-1_ is redox active and is an iron oxidase. UV-Vis spectra of Cyc2_PV-1_ after enrichment chromatography and cleavage of the His-tag. (A) Cyc2_PV-1_ as purified was oxidized (black). Addition of potassium hexacyanoferrate (III) showed no changes to the heme spectra, confirming the oxidized state (gray). Sodium dithionite reduced Cyc2_PV-1_ (red), as evidenced by the shift in the Soret peak and the appearance of the alpha and beta peaks (black arrows). (B) Cyc2_PV-1_ as purified (black). Fe(II) reduced Cyc2_PV-1_ (red), shown by the shift in the Soret peak and the appearance of the same alpha and beta peaks as the dithionite-reduced spectra.

To perform the iron oxidation assay, we used Fe(II) with citrate to chelate the Fe(III) product, preventing formation of iron oxyhydroxide precipitates that would interfere with the UV-Vis assay. Citrate is a weak Fe(II) ligand (stability constant log *K* = 3.20; https://www.nist.gov/srd/nist46), readily releasing Fe^2+^. We calculated that ∼20% of the Fe(II) is Fe^2+^ in our assay solution (Supplementary Table 2; (61)), so Fe^2+^ is available as a substrate. Reaction between Fe(II) and Cyc2_PV-1_ showed the same shift of the Soret peak and appearance of the same alpha and beta peaks as reduction by sodium dithionite (Fig. 5B). These spectral changes show Cyc2_PV-1_ can accept electrons from Fe(II), demonstrating that Cyc2_PV-1_ is an iron oxidase.

### Cyc2_PV-1_ has a redox potential of 208 ± 20 mV

To determine the redox potential of Cyc2_PV-1_, we used a modified Massey method protocol developed for low-yield heme proteins (62–64). Cyc2_PV-1_ was titrated against 2,6-dichlorophenolindophenol (DCPIP) with a known redox potential of +217 mV (63). By plotting the Nernst equation-transformed ratios of oxidized and reduced forms of both DCPIP and Cyc2_PV-1_ from four independent experiments (Fig. S4), we calculated the redox potential of Cyc2_PV-1_ as 208 ± 20 mV (Fig. 6).

**Figure 6.**
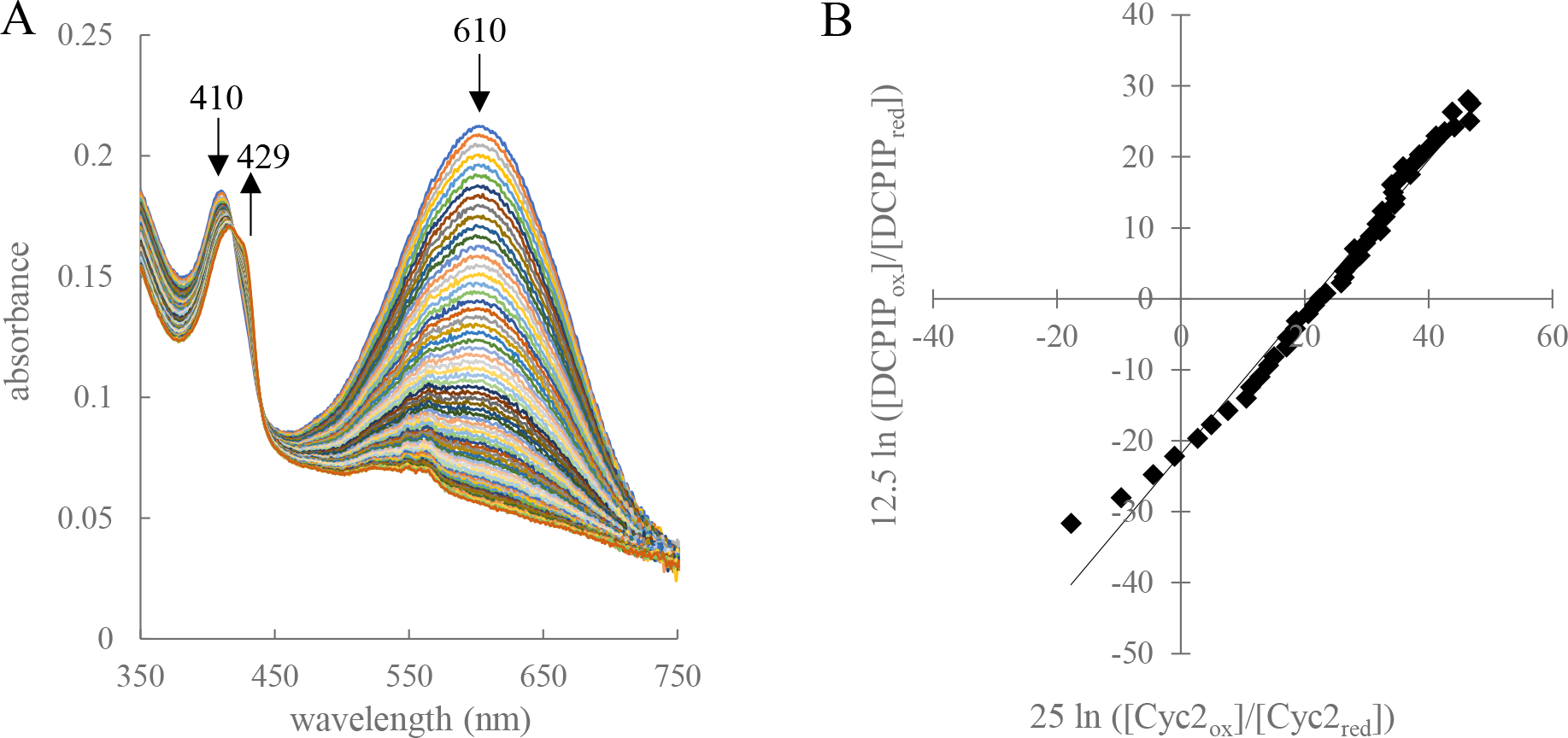
Cyc2_PV-1_ redox potential measurements shown by representative UV-Vis spectra and data plotting. (A) Spectroscopic changes observed during the determination of the reduction potential of Cyc2_PV-1_ using the dye DCPIP. UV-Vis spectra were recorded every 15 sec during a xanthine oxidase-catalyzed reductive titration with xanthine, DCPIP, and Cyc2_PV-1_. (B) The ratios of the reduced and oxidized forms of Cyc2_PV-1_ and DCPIP were plotted and used to determine the redox potential of Cyc2_PV-1_.

Our results show a redox interaction between Cyc2_PV-1_ and Fe(II) (Fig. 5), and the calculated redox potential of Cyc2_PV-1_ (Fig. 6) puts it between the redox potentials of Fe(III)/Fe(II) (pH 7) and O_2_/H_2_O (Fig. 7), as would be expected for a neutrophilic iron-oxidizer. In contrast, the redox potential for Cyc2 from *A. ferrooxidans* was measured at 560 mV (27) which is consistent with the higher Fe(III)/Fe(II) redox potential at low pH. These differences suggest Cyc2 can be tuned to different redox potentials suitable for different FeOB environments.

**Figure 7.**
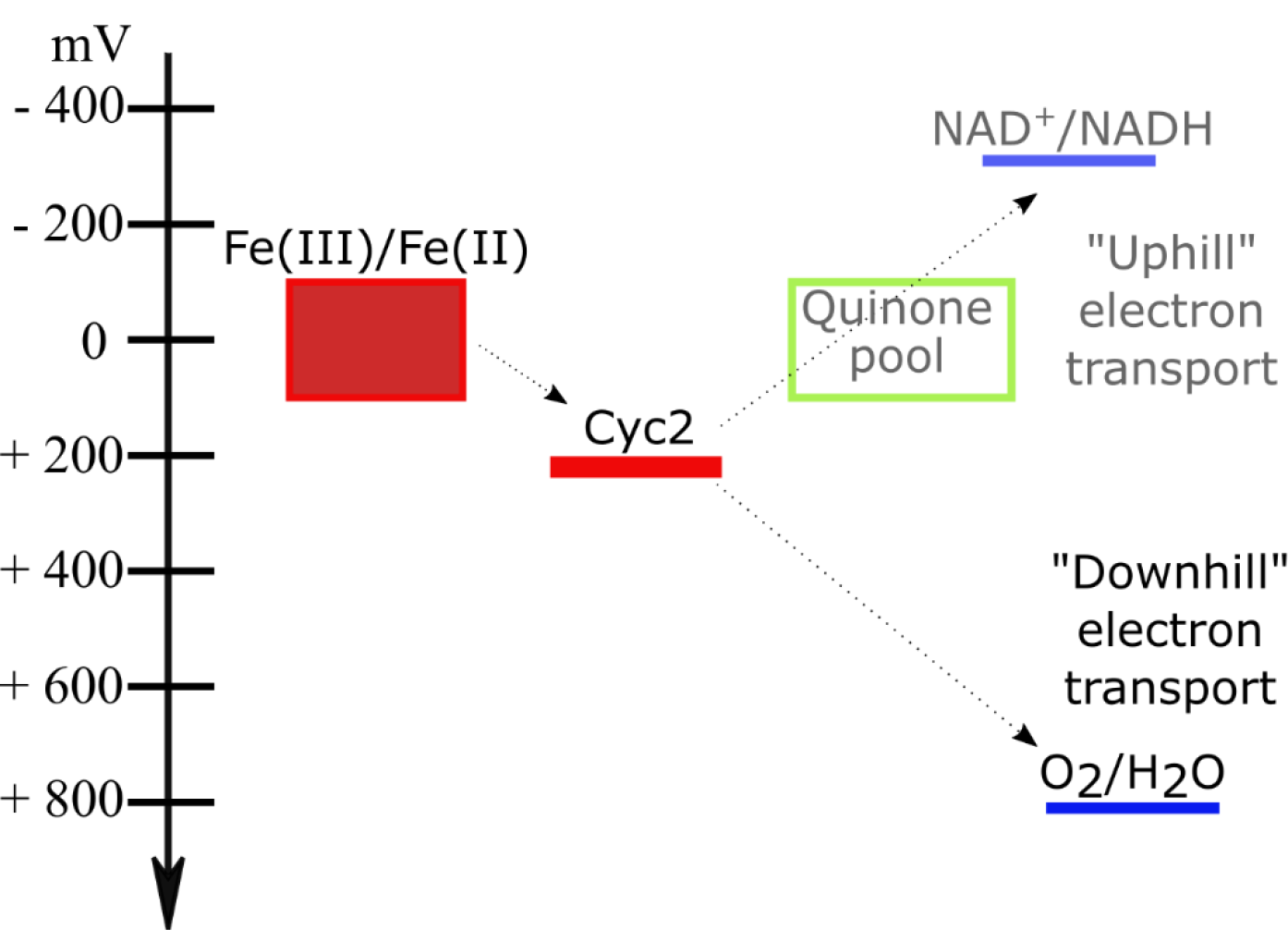
Schematic of reduction potentials showing how Cyc2_PV-1_ fits into the neutrophilic electron transport pathway. Boxes represent the range of reported values: for both the Fe(III)/Fe(II) couple (Fe^2+^/ferrihydrite couple, as predicted from FeOB mineralogy) and menaquinones and ubiquinones, this ranges -110 to +110 mV. Much remains unknown in the “downhill” electron transport to the terminal electron acceptor, oxygen, but likely involves other cytochromes (11, 34).

### Differentiating the structure of Cyc2 from other iron oxidoreductases

In total, these results confirm that Cyc2 is an outer membrane iron oxidase. While Cyc2 is unique in that the cytochrome and porin are fused, its overall structure is reminiscent of other porin cytochrome complexes that play roles in iron cycling, particularly MtrCAB/OmcA and MtoAB/PioAB (30–32, 65–67). These complexes include a 26-strand beta barrel that accommodates insertion of a decaheme cytochrome, which spans the outer membrane and may contact other extracellular decaheme proteins to conduct extracellular electron transfer (Fig. 8). In contrast, Cyc2 is predicted to have a single heme and a smaller barrel (16 strands). For organisms that eke out a living from iron oxidation, the single heme and smaller size of Cyc2 means it requires fewer resources to produce, making it an attractive alternative to larger porin cytochrome complexes (68).

**Figure 8.**
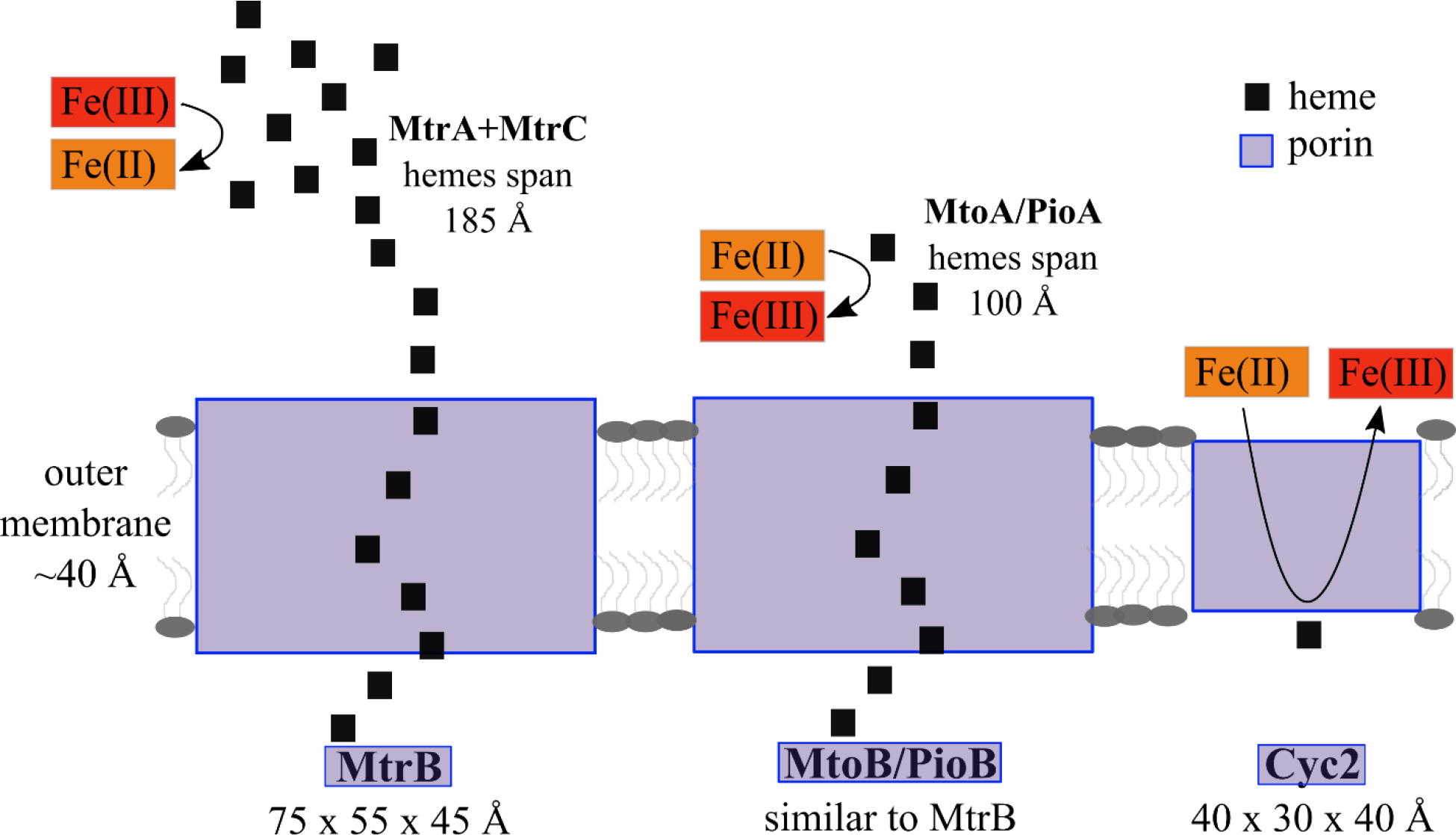
Cyc2 is different than other iron oxidoreductases. The MtrCAB complex is a 26-stranded porin with hemes spanning a range of 185 Å (66). The PioAB/MtoAB complex is likely similarly sized to MtrAB, with hemes spanning up to ∼100 Å based on modeling (32). Cyc2 is a smaller porin of 16-strands and possesses only a single heme. Due to its smaller size and placement of single heme, Fe^2+^ would have to enter the pore of Cyc2.

If the pore size of Cyc2 is similar to the structurally homologous phosphate porins, the internal diameter is expected to be ∼3.5 Å (46). A model of the Cyc2_PV-1_ cytochrome domain is ∼20 x 20 x 10 Å (Fig. S5), which would fit as a plug within the periplasmic opening of the barrel, but not span the outer membrane through the porin, nor fully reside within the middle of the pore. Given our structural predictions, the cytochrome would be on the periplasmic side of Cyc2, which would allow transfer of electrons to a periplasmic component of the electron transport chain. While long range electron transport is possible, the rate of electron transport decreases exponentially with distance (69). The hemes within MtrA are 3.9-6.5 Å apart (66), and similar configurations are found in other cytochromes as well, suggesting this distance is optimal for efficient electron transport. In our model, Fe^2+^ would need to enter the Cyc2 pore to some extent, possibly with the help of a chaperone or ligand that could also escort Fe^3+^ from the pore before formation of Fe(III) oxyhydroxides that would detrimentally encrust the protein. The requirement for iron to enter the pore would suggest that Cyc2 is an oxidase of dissolved Fe^2+^, distinguishing it from a multiheme iron oxidase like MtoA that could directly conduct electrons from a solid surface.

### Recognizing cyc2/Cyc2 in other organisms

In the short time since the first comprehensive Cyc2 phylogenetic tree was published (7), many more *cyc2* homologs have been sequenced. To explore how these new sequences fit into our understanding of Cyc2 phylogeny, we updated the Cyc2 tree with sequences from databases that met the criteria of our in-house pipeline--specifically a minimum length (365 amino acids), presence of a heme-binding site, and low blastp similarity cut-off (1E10^-5^). Even with the substantial increase in Cyc2 homologs, the topology of the tree remained the same, with strong support for three main clusters of Cyc2 sequences (Fig. 9; figshare File 2: <u>10.6084/m9.figshare.c.5390285</u>).

**Figure 9.**
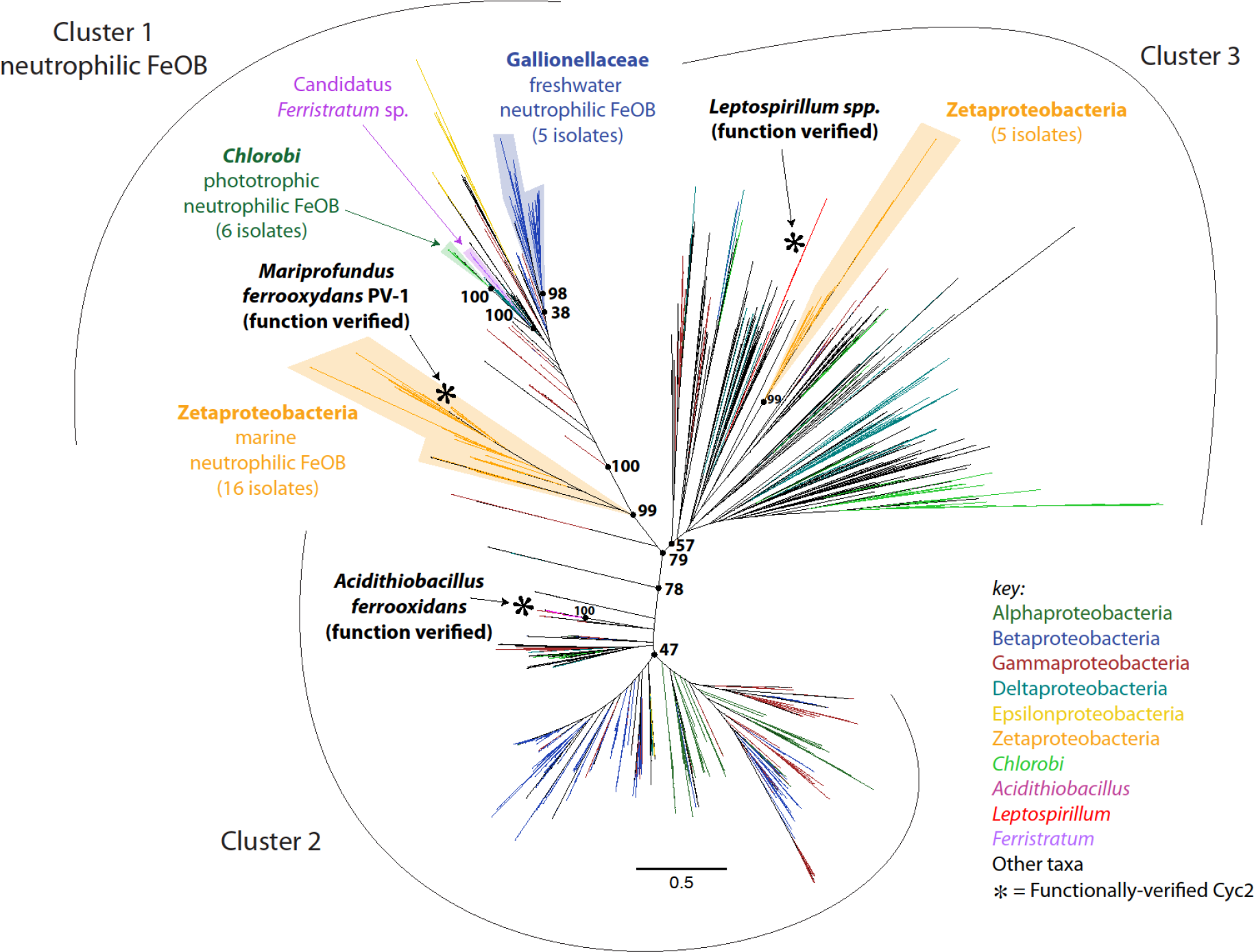
Updated Cyc2 maximum likelihood phylogenetic tree (1,593 sequences total). Sequences come from isolates, enrichments, single amplified genomes, and metagenome assembled genomes, with few replicates. The three Cyc2 clusters continue to be supported (79% bootstrap support) and are confirmed with HMMs from FeGenie. Functionally-characterized Cyc2 are shown with asterisks, with one from each cluster. Known neutrophilic Fe(II)-oxidizing clades are labelled with the number of distinct isolates represented by each. Full rectangular version of tree with leaf labels in figshare File 2: <u>10.6084/m9.figshare.c.5390285</u>.

We have historically relied on our in-house pipeline for identifying Cyc2 homologs because the Cyc2 sequence is dominated by a porin domain that characteristically features low amino acid conservation (70). In contrast, the short cytochrome domain is the most conserved region (Fig. S6, Fig. S7), and a consensus sequence derived from all homologs includes a AXPXFAR[Q/K][T/Y] motif located 5 amino acids upstream of the CXXCH heme binding site, and a PXL motif 4 amino acids downstream of the CXXCH (Fig. S8). The PXL motif can be found in many other cytochromes, such as MtoD (71) and Cyc1 in *A. ferrooxidans* (CYC41 in (72)); the proline and lysine appear to help stabilize the heme (72). In contrast, the AXPXFAR[Q/K][T/Y] motif is unique to Cyc2, and therefore can be used to identify Cyc2-like cytochromes.

Recently, we developed hidden Markov models (HMMs) to identify Cyc2 sequences. These HMMs are part of the FeGenie program (73), which identifies and assigns putative Cyc2 homologs to one of the three phylogenetic clusters, depending on which HMM is the highest-scoring match. The HMMs were built from representative sequences of the phylogenetic tree presented by McAllister *et al*. (7), and each phylogenetic cluster is represented by a unique HMM. An alignment of sequences from each cluster show some differences in the consensus motif between clusters (Fig. S8). The FeGenie algorithm confirms presence of a heme-binding motif and excludes sequences shorter than 365 amino acids, but users should confirm the presence of a porin in FeGenie-identified Cyc2 homologs using secondary structure prediction software (e.g. HHPRED). We confirmed the FeGenie Cyc2 HMM cluster classification matched the location of the sequences in the updated Cyc2 tree, with only one exception (figshare File 2: <u>10.6084/m9.figshare.c.5390285</u>). The Cyc2 HMMs have been validated and are the recommended tool for finding new Cyc2 homologs (73).

### Interpreting Cyc2 sequences: towards the discovery of new iron-oxidizers

We now have biochemical evidence for iron oxidation function by three diverse Cyc2 representatives that each fall within one of the three main clusters. It is tempting to now attribute iron oxidation function to the various Cyc2 homologs found in organisms across the tree (Fig. 9), as well as new homologs found using the HMM-based strategy above. However, the degree of confidence depends heavily on context, particularly how closely related the homolog is to a characterized Cyc2. New sequences can be added to the Cyc2 phylogenetic tree using the full alignment file (figshare File 3 and 4: <u>10.6084/m9.figshare.c.5390285</u>). In some cases, the new sequence may fall within a well-supported clade of known FeOB with good evidence for Cyc2 function, which will give confidence in its predicted role in iron oxidation. However, the many long branches on the Cyc2 tree indicate a large degree of divergence, notably in Cluster 3. The functionally characterized *Leptospirillum* Cyt_572_ has an especially long branch, and also has an unusual heme prosthetic group (74), both of which could mean that Cyt_572_ is not necessarily representative of Cluster 3. In all, the function of *cyc2* homologs within Cluster 3 may be considered most uncertain, requiring further verification.

In contrast, Cluster 1 has shorter branches, indicating that the sequences are more closely related to one another and to the biochemically tested Cyc2_PV-1_. In fact, many of the sequences (∼ 55%) within this cluster are from well-characterized neutrophilic FeOB, i.e. the Gallionellaceae, Zetaproteobacteria or Chlorobi, including all known iron-oxidizing isolates of those taxa. In addition, previous work has helped build a case for prevalent Cluster 1 Cyc2 function. Cluster 1 FeOB *cyc2* are highly expressed in iron-oxidizing environments, including Gallionellaceae *cyc2* in the Rifle alluvial aquifer (5) and Zetaproteobacteria *cyc2* in marine iron mats in three hydrothermal systems (7). This hints at their importance in iron oxidation, which is further supported by expression that corresponds to iron-oxidizing conditions stimulated in microcosms (7). Building on this, here we show that a Cluster 1 Cyc2 oxidizes Fe(II) and has an appropriate redox potential. Altogether, this gives a higher degree of confidence in generalizing the iron oxidation function across Cluster 1 Cyc2.

The Cyc2 phylogenetic tree shows that Cyc2 sequences are very diverse, with many homologs that are distantly related to Cyc2_PV-1_, *A. ferrooxidans* Cyc2, and Cyt_572_, notably much of Clusters 2 and 3. Thus, there is still much work to be done to verify Cyc2 function in both known FeOB and organisms not known to oxidize iron. For known FeOB, we are building the capacity for genetics-based validation (75). But the vast majority of *cyc2*/Cyc2 sequences come from genomes of uncultured organisms. In these cases, how can we test the function? We recently discovered Cluster 1 *cyc2* sequences in metagenome-assembled genomes of *Candidatus* Ferristratum spp. from the DTB120 phylum (Fig. 9), found in deeper portions of hydrothermal iron microbial mats inhabited by well-characterized Zetaproteobacteria FeOB. To test whether the new *Ca.* Ferristratum *cyc2* is connected to iron oxidation, we added Fe(II) to iron mat sample microcosms and analyzed a time course of transcriptomes. The expression of *cyc2* and the nitrate reduction gene *narG* from *Ca.* Ferristratum increased concurrently in response to Fe(II) addition, suggesting this group of organisms couples iron oxidation to nitrate reduction (10). This approach can guide the exploration of the diversity, distribution and roles of iron-oxidizing bacteria in environmental systems. If many other Cyc2 homologs prove to be iron oxidases, microbial iron oxidation may be more widespread than we currently recognize.

## Experimental Procedures

### Structural modeling

Signal peptides were predicted using SignalP (http://www.cbs.dtu.dk/services/SignalP/) (40) and secondary elements were predicted using PSIPRED (http://bioinf.cs.ucl.ac.uk/psipred/) (42, 43). For identification of structural homologs to Cyc2, we uploaded sequences to the HHpred tool (https://toolkit.tuebingen.mpg.de/tools/hhpred) and searched against the PDB_mmCIF70_14_Oct database, available as part of the Max Planck Institute Bioinformatics Toolkit (44, 45, 48). Structural modeling was carried out by HHpred/MODELLER (48, 51), I-TASSER (49), and Phyre2 (50). The structural models from all three platforms were found to be in close agreement.

### Cloning and heterologous expression of Cyc2_PV-1_

The *cyc2* gene of *Mariprofundus ferrooxydans* PV-1 (accession no. AKN78226) was codon-optimized for expression in *Escherichia coli* and synthesized by Genscript (Piscataway, NJ, USA) with a C-terminal *Strep*-tag II (WSHPQFEK) (76). The native PV-1 signal sequence was replaced with the signal sequence of an *E. coli* outer membrane porin, *ompA* (WP_063091005 amino acids 1-21). The *cyc2* gene was cloned into the EcoRI/ HindIII sites of the pMal-p4X plasmid (with the *malE* gene removed). This *cyc2*-containing plasmid was co-transformed into *E. coli* C41(DE3) (Lucigen) with pEC86*, a plasmid containing the cytochrome c maturation (*ccmABCDEFGH*) genes under a constitutive promoter, to ensure heme insertion into Cyc2_PV-1_ (77). The pEC86 used here (pEC86*) was a spontaneous mutant found to possess a frame-shift and early stop codon in the *ccmA* gene. For unknown reasons that are beyond the scope of this manuscript, this plasmid was not detrimental to yield and beneficial to detection of Cyc2_PV-1_ in our system, and was used in all expression studies.

The porin-only construct was created by removing the bases corresponding to the cytochrome domain from the plasmid via site-directed mutagenesis. The 8xHis-tag, TEV protease cleavage site, and linker region were inserted into the relevant constructs also by site-directed mutagenesis (NEB Q5-based kit). Optimized DNA sequence is provided in figshare File 1 (<u>10.6084/m9.figshare.c.5390285</u>). Plasmids were propagated in *E. coli* NEB5α competent cells, and clones were sequenced after manipulation by Sanger sequencing at the University of Delaware Sequencing and Genotyping Center prior to being transformed into *E. coli* C41(DE3)/pEC86*.

For expression of *cyc2*, *E. coli* was grown aerobically at 37 °C (with shaking at 200 RPM) in Lysogeny Broth (LB), with ampicillin (100 µg/mL) for propagation of the pMal-p4X based-plasmids, and chloramphenicol (30 µg/mL) for the pEC86* plasmid. LB was routinely supplemented with 2 mM MgCl2, trace vitamins and minerals (78), and ferric citrate. After reaching an OD_600_ of ∼0.5, cultures were induced with 0.1 mM isopropyl β-D-1-thiogalactopyranoside (IPTG) for de-repression of the *lac* operon; induction proceeded for 3-4 hours at 30 °C with shaking at ∼200 RPM. We evaluated the use of 5-aminolevulic acid for improving expression (79, 80) but found no benefit.

### Purification of Cyc2_PV-1_ and assays

Cells were harvested by centrifugation (6000 x *g*), 15 min at 4° C and the pellet stored at -20° C until lysed. Cells were lysed by a passage through the Avestin C5 Emulsiflex homogenizer at 10000-15000 psi in 20 mM Tris pH 8 with protease inhibitors and 10 µg/mL DNAse. After lysis, cellular debris was removed by centrifugation at 15000 x *g*, 20 min at 4 °C. Total membranes were harvested by ultracentrifugation at 100000 x *g*, 1 hr, 4 °C in a Beckman Coulter SW28 or SW32 rotor. Total membranes were resuspended in 20 mM Tris pH 8/1.5 M sucrose/1% DDM. The membranes were solubilized by rotating at 4 °C overnight. Solubilized membranes were mixed 1:1 with binding buffer (20 mM Tris pH 8/1 M NaCl/20 mM imidazole) and loaded onto a HisPur Ni-NTA (Thermo Scientific) column equilibrated with 20 mM Tris pH 8/750 mM sucrose/500 mM NaCl/10 mM imidazole/0.2% DDM (equilibration buffer). The column flow-through was collected and the column washed with 10 column volumes of equilibration buffer. After washing, bound protein was eluted with 300 mM imidazole in equilibration buffer.

For the redox assays, eluted protein containing Cyc2_PV-1_ was dialyzed against equilibration buffer to remove the high concentration of imidazole. Then the His-tag was cleaved by incubation with TEV protease (NEB) for 18 h at room temperature. Following cleavage, the fraction was re-loaded onto an equilibrated Ni-NTA column and tag-free Cyc2_PV-1_ was not retained by the column while TEV protease was. The tag-free Cyc2_PV-1_ was dialyzed against assay buffer (20 mM MES pH 6.3/300 mM NaCl/500 mM sucrose/0.1% DDM) to adjust the buffer conditions for the assay. After dialysis, the protein was concentrated using a 30 kDa MWCO spin concentrator (Amicon Ultra). Cyc2_PV-1_ was degassed and taken into the anaerobic chamber (Coy 5% H_2_/95% N_2_ atmosphere) and diluted into anoxic assay buffer. Diluted Cyc2_PV-1_ was sealed in a masked, quartz microcuvette with septum lid (Starna Cells), then removed from the chamber for UV-Vis readings on a NanoDrop OneC Spectrophotometer (Thermo Scientific). The microcuvette was returned to the anaerobic chamber, and 2 mM ferrous citrate was added to the cuvette, before it was resealed and taken back out to the spectrophotometer. Spectra from 200-800 nm were collected every min for ∼30 minutes. The reduced Cyc2_PV-1_ spectra (Fig. 5A) were obtained by adding a final concentration of 5 mM sodium dithionite to Cyc2_PV-1_ in assay buffer. The oxidized Cyc2_PV-1_ spectra (Fig. 5A) were obtained by adding a final concentration of 100 µM potassium hexacyanoferrate (III) to Cyc2_PV-1_ in assay buffer.

The redox potential was measured by a modified Massey method (62–64). Cyc2_PV-1_ was purified as above, then concentrated and buffer-exchanged into 20 mM Tris pH 8/300 mM NaCl/300 mM sucrose/0.02% DDM using the Amicon spin concentrator. Cyc2_PV-1_ was degassed and taken into the anaerobic chamber and diluted into anoxic assay buffer. Diluted Cyc2_PV-1_ was sealed in a masked, quartz microcuvette with septum lid, then removed from the chamber for UV-Vis readings on an Agilent Cary3500 spectrophotometer. Cyc2_PV-1_ was taken back in the chamber for addition of the reference dye, 2,6-dichlorophenolindophenol (*E*_m_ = 217 mV) and 500 µM xanthine (both from Sigma-Aldrich), then sealed and removed to take UV-Vis readings on the spectrophotometer. The reaction was initiated at the spectrophotometer by adding anoxic xanthine oxidase (microbial; 1-3 µg/mL; Sigma-Aldrich) via syringe and needle through the septum lid. Readings of 350-800 nm were collected every 15 sec. for ∼1 hour. The absorbance change for the reduced Cyc2_PV-1_ Soret maximum was monitored at 429 nm, at which there was negligible contribution from DCPIP, and the absorbance change for the dye peak was monitored at 610 nm, at which there was negligible contribution from Cyc2_PV-1_. These changes in absorbance were transformed by the Nernst equation and plotted: the one-electron reduction of Cyc2_PV-1_ with [25 mV ln (Cyc2_red_/Cyc2_ox_)] and the two-electron reduction of DCPIP with [12.5 mV ln (DCPIP_red_/DCPIP_ox_)]. Data points where the reduced/oxidized ratio was greater than 10 or less than 0.065 were excluded from the analysis. The y-intercept of a line fit to the data represents the difference in potential between the heme and the known potential of DCPIP. The reduction potential of Cyc2_PV-1_ was calculated based on four independent titrations. Bovine heart cytochrome c (Sigma-Aldrich) was used as a positive control to optimize experimental conditions.

### SDS-PAGE, western and heme staining

Whole cells to be analyzed by SDS-PAGE were lysed by resuspension of a cell pellet in 5X SDS running buffer (125 mM Tris, 1.25 M glycine, 0.5% SDS, pH 8.3) and passing the resuspension through a 27.5 gauge needle 10 times. Samples were incubated 1:1 in 2X loading buffer (250 mM Tris, 2% SDS, 30% glycerol, 0.002% bromophenol blue) prior to loading on a 15% Tris-buffered polyacrylamide resolving gel, topped with a 5% polyacrylamide stacking gel, and electrophoresed according to the method of Laemmli (81). Samples were electrophoresed at 70 V for 30-45 min, then 125 V for 45-60 min. The gel was then either Coomassie-stained (1 g Coomassie Brilliant Blue Stain in 10% acetic acid, 40% ethanol), or heme-stained. For heme-staining (82), 0.1 g o-dianisidine was dissolved in 90 mL water. Immediately prior to the start of staining, 10 mL sodium citrate pH 4.4 and 200 µL hydrogen peroxide were added to the o-dianisidine solution. Heme-containing proteins in the gel turned brown after 15 min - 2 hours incubation in the heme stain at room temperature. For Western blot and another version of the heme stain, the proteins from the SDS-PAGE gel were transferred to a PVDF membrane at 30 V for 16 h (4°C) in transfer buffer (25 mM Tris, 192 mM glycine, 20% methanol, pH 7.2). Heme peroxidase activity was assessed by washing the membrane with TBST buffer (20 mM Tris, 137 mM NaCl, 0.1% Tween-20, pH 7.6), and incubating for 15-30 minutes with Pierce ECL luminol and substrate (83), before imaging on a Typhoon FLA 9500 (GE Healthcare Life Sciences) or iBright FL1500 system (Invitrogen). For *Strep*-tag II detection, the PVDF membrane was blocked for one hour with 1% bovine serum albumin (BSA) in TBST buffer, rinsed with TBST, and incubated for one hour with Precision Protein Streptactin-HRP conjugate (1:60,000 dilution, Biorad). The membrane was then washed with four 5-minute washes in TBST, then incubated with Pierce ECL luminol and substrate for 5 minutes before imaging on the Typhoon FLA 9500 or iBright system.

### Separation of inner and outer membrane layers

Cyc2_PV-1_ was expressed in *E. coli* C41(DE3)/pEC86* as described and cells were lysed on the Emulsiflex as above. After lysis, cellular debris was removed by centrifugation at 15000 x *g*, 20 min. at 4 °C. A sucrose gradient was set up with 7 mL 1.5 M sucrose as the bottom layer, 7 mL 0.5 M sucrose as the middle layer, and 15 mL lysed cells as the top layer (58, 59). The gradient was ultracentrifuged at 100000 x *g*, 18 hr at 4 °C in a Beckman Coulter SW28 or SW32 rotor. Following ultracentrifugation, the outer membrane could be found at the bottom of the tube, and the inner membrane localized to the interface between the 1.5 M and 0.5 M sucrose layers. Cytoplasmic proteins remained above the 0.5 M sucrose layer.

### Mass spectrometry

#### Sample preparation

Gel bands of interest were excised for analysis. The enzymatic digestion procedure was performed with trypsin (Promega) at 37 °C as previously described (84) and included reduction/alkylation with dithiothreitol (BioRad) and iodoacetamide (Sigma), respectively. Subsequently, samples were desalted and concentrated using µC18 Ziptips (Millipore) per manufacturer’s instructions, then dried with a SpeedVac vacuum concentrator (Thermo Fisher) and resuspended in 24 µL 2% acetonitrile (ACN), 0.1% formic acid (FA).

#### LC-MS/MS data acquisition

Liquid chromatography-tandem mass spectrometry (LC-MS/MS) was performed on a TripleTOF 6600 (Sciex) coupled to an Eksigent nanoLC 425 (Sciex) operating in microLC flow mode. Twelve microliters of each sample were injected onto a ChromXP C18CL column (3 µm, 120 Å, 150 mm x 0.3 mm, Sciex). Gradient elution was performed with mobile phase A (0.1% formic acid in water, Fisher) and B (0.1% formic acid in ACN, Fisher) at a flow rate of 5 µL/min. A program of 3% B to 35% B in 5 min, 35% B to 80% B in 1 min, at 80% B for 2 min was used to elute peptides. The eluate was ionized with a dual spray source. The mass spectrometer was operated in positive ion mode with a full MS1 scan experiment in a mass range of 400-1250 m/z with a scan time of 250 ms followed by MS/MS experiments in the mass range of 100-1500 m/z with a scan time of 50 ms. The top 30 precursor ions were selected for fragmentation.

#### Database search

LC-MS/MS data were searched against a database of all *E. coli* K-12 protein sequences (available at: ftp.ncbi.-nih.gov), modified to include Cyc2_PV-1_, with ProteinPilot v5.1 (Sciex) using the Paragon algorithm. Search parameters were specified as following: iodoacetamide cysteine modification, trypsin enzyme, gel ID special factors. Protein identifications with local FDR 1%, and at least 2 high confidence unique peptides (99% confidence) were considered.

### Cyc2 phylogeny and amino acid identity calculations

The Cyc2 phylogenetic tree was constructed following the same methods as published previously (7). The database of 634 full-length dereplicated Cyc2 sequences was updated by adding new sequences identified by blastp (85) against the NCBI and IMG databases (maximum e-value of 1*10^-5^) using one query from each of the three Cyc2 clusters. Resulting sequences were dereplicated using cd-hit at 100% identity over 90% of the length of the protein (86). The database was further screened to remove sequences either shorter than 365 amino acids, lacking the cytochrome c binding motif (CXXCH) or upstream PXFAR[Q/K][T/Y] motif, and several from highly sampled genera (*Burkholderia*, *Ralstonia*, and *Xanthomonas*). This resulted in a total of 1,593 full-length sequences. To generate the tree, these sequences were aligned using MUSCLE (87) (figshare File 3: <u>10.6084/m9.figshare.c.5390285</u>), the alignment was masked to remove positions with greater than 30% gaps (figshare File 4: <u>10.6084/m9.figshare.c.5390285</u>), and a maximum likelihood phylogenetic tree was built using RAxML (276 alignment columns, 300 bootstraps, CAT model of rate heterogeneity, JTT amino acid substitution model) (88). The resulting phylogenetic tree was colored and names customized using the Iroki program (89). Cyc2 cluster assignments were confirmed by running all sequences through the FeGenie program (73). Amino acid identity (AAI) values were calculated from the pairwise alignments prior to masking.

### HMM development and calibration

Representative sequences from each Cyc2 phylogenetic cluster were aligned using MUSCLE (87), then manually curated. These alignments were then used to construct HMMs with HMMER (90). The HMMs were calibrated by querying each HMM against NCBI’s non-redundant (nr) protein database. The results of each search were visually inspected to identify the optimum bit score cutoff for each HMM. Due to the poor conservation of the porin domain of Cyc2, many Cyc2 homologs were identified at low bit scores; these low-scoring homologs were confirmed by 1) visual inspection of the sequence at the N-terminus to confirm presence of the conserved cytochrome motif and 2) secondary-structure modelling, using HHPRED (48). Occasionally, false positives were identified at similar bit score values to many true positives (e.g. porins without heme-binding motifs). To exclude these types of false positives, any protein matching a Cyc2 HMM without a heme-binding motif is automatically excluded from the results through a built-in heme-detecting function. Additionally, because the vast majority of confirmed Cyc2 homologs are at least 400 residues in length, FeGenie is programmed to exclude any match to the Cyc2 HMMs shorter than 365 residues.

## Acknowledgments

This research was funded by the National Science Foundation (EAR-1151682, BIO-1817651) and the Office of Naval Research (N00014-17-1-2640). We thank the Delaware Biotechnology Institute (DBI) for core instrumentation. Support from the UD CBCB Bioinformatics Core Facility and use of the BIOMIX computer cluster was made possible through funding from Delaware INBRE (NIH NIGMS P20 GM103446), the State of Delaware, and the Delaware Biotechnology Institute. We acknowledge Shawn Polson for assistance in bioinformatic analyses, the UD Sequencing and Genotyping Center for Sanger Sequencing, and Leila Choe and Kelvin Lee of the DBI Proteomics and Mass Spectrometry Core for performing the mass spectrometry. We thank Dianne Newman and Lina Bird for the pEC86 plasmid. We are grateful to the Rozovsky lab members for assistance with experiments, to Hans Carlson and Tom Hanson for advice, and to Nanqing Zhou and Rene Hoover for helpful comments on the manuscript. The authors declare no conflict of interest.

## Supplemental files

**Figure S1.** PSIPRED prediction of secondary structure in Cyc2_PV-1_, the Cyc2 sequence from *M. ferrooxydans* PV-1 (D. W. A. Buchan, D. T. Jones, Nuc Acids Res, 2019, https://doi.org/10.1093/nar/gkz297).

**Figure S2.** Constructs and expression of Cyc2_PV-1_. Cyc2_PV-1_ is marked with a red arrowhead, protein ladder is labeled with an M (Spectra Broad Range on heme and Coomassie, WesternC on α-*Strep*-tag Western blot), and relevant band sizes are labeled in kDa. (A) Schematic of gene construct for expression and representative stained SDS-PAGE gels showing Cyc2_PV-1_ expression in *E. coli*: 1) uninduced, 2) induced, 3) lysed and induced. Smaller bands visible on *Strep*-tag Western blots are non-specific. (B) Heme-stained gel of fractions during His-tag purification. Cyc2_PV-1_ migrates at its expected molecular weight in the diluted total membranes (input) and flow-though (FT). After elution, Cyc2_PV-1_ migrates in a high-molecular weight complex (1 - 300 µM imidazole, 2 - after dialysis to remove imidazole). After TEV protease cleavage of the His-tag, Cyc2_PV-1_ does not interact with the Ni-NTA column (3 - flow-through, 4- imidazole elution). (C) Uncropped gel corresponding to Figure 2D. See lane labels to the right of image.

**Figure S3.** Uncropped gels. (A) Coomassie-stained gel corresponding to Fig. 3A. CP1, CP2 - cytoplasmic proteins, IM - inner membranes, OM - outer membranes, M - Spectra Broad Range protein ladder with relevant bands in kDa. (B) Gels corresponding to samples in Fig. 4. Red arrow indicates full-length Cyc2_PV-1_ and blue arrow indicates porin-only. Lanes: 1-empty vector, 2-porin lysed supernatant, 3-porin ultracentrifuged supernatant, 4-porin total membranes, 5-porin cytoplasmic proteins, 6-porin inner membranes, 7-porin outer membranes, 8-Spectra Broad Range or WesternC protein ladder, 9- Cyc2_PV-1_ lysed supernatant, 10- Cyc2_PV-1_ ultracentrifuged supernatant, 11- Cyc2_PV-1_ total membranes, 12- Cyc2_PV-1_ cytoplasmic proteins, 13- Cyc2_PV-1_ inner membranes, 14- Cyc2_PV-1_ outer membranes.

**Figure S4.** Four independent redox titration reactions with Cyc2_PV-1_ and DCPIP.

**Figure S5.** Three views of the modeled cytochrome domain of Cyc2_PV-1_. The view on the right is rotated 90 ° away from the viewer compared to the view in the center. Hydrophobic residues are gray and polar residues are red. Heme (not pictured) is covalently attached to cysteine residues (yellow) and coordinated by histidine (blue). Model generated using MODELLER (B. Webb, A. Sali, Curr Prot Bioinf, 54: 5.6.1-5.6.37, 2016, https://doi: 10.1002/cpbi.3).

**Figure S6.** Full alignment of Cyc2 from representative neutrophilic and acidophilic FeOB. Orange line indicates the conserved cytochrome region.

**Figure S7.** Histograms of pairwise amino acid identity of the (A) full length Cyc2 sequences, (B) porin portion, and (C) cytochrome portion (n=156). The cytochrome portion is more highly conserved than the porin. (D) Amino acid identities (AAI) of full length Cyc2 sequences from FeOB and *Tenderia electrophaga*. AAI to biochemically characterized Cyc2 are shown in bold. Note that organisms from Cluster 1 e.g. neutrophilic FeOB Zetaproteobacteria, Gallionellaceae, and *Chlorobi* are most similar to one another.

**Figure S8.** Comparison of motifs found in the conserved cytochrome domain of Cyc2. The sequence logo labeled “All” is built from 1593 homologs. Each of the Cluster logos are built from all sequences in each cluster (334 in Cluster 1, 858 in Cluster 2, and 401 in Cluster 3).

**Table S1.** Unique peptides detected by tandem MS/MS that matched to Cyc2_PV-1_ with 99% confidence

**Table S2.** Relevant Fe(II) and citrate speciation in the iron oxidase assay buffer from Visual MINTEQ calculation (https://vminteq.lwr.kth.se/).

**figshare files available at** <u>10.6084/m9.figshare.c.5390285</u>:

File 1: Full-length Cyc2_PV-1_ sequences. Full-length nucleotide and amino acid sequence for Cyc2_PV-1_ expression construct used in purification.

File 2: Full rectangular Cyc2 phylogenetic tree. This Cyc2 maximum likelihood phylogenetic tree shows all bootstraps and leaf labels. FeGenie cluster calls are shown as different colored dots to the left of the leaf labels.

File 3: Full Cyc2 alignment file. MUSCLE alignment file of all 1,593 sequences used in the Cyc2 phylogenetic tree. All amino acid positions in this alignment have been preserved.

File 4: Masked Cyc2 alignment file. MUSCLE alignment file of all 1,593 sequences used in the Cyc2 phylogenetic tree. Positions with greater than 30% gaps were removed.

## References

1. Borch T, Kretzschmar R, Kappler A, Cappellen PV, Ginder-Vogel M, Voegelin A, Campbell K. 2010. Biogeochemical redox processes and their impact on contaminant dynamics. Environ Sci Technol 44:15–23.

2. Emerson D, Fleming EJ, McBeth JM. 2010. Iron-oxidizing bacteria: an environmental and genomic perspective. Annu Rev Microbiol 64:561–583.

3. Emerson D, De Vet W. 2015. The role of FeOB in engineered water ecosystems: A review. Journal - American Water Works Association 107:E47–E57.

4. Kappler A, Emerson D, Gralnick JA, Roden EE, Muehe EM. 2015. Geomicrobiology of iron., p. 343–399. In Ehrlich’s Geomicrobiology, 6th ed. Boca Raton.

5. Jewell TNM, Karaoz U, Brodie EL, Williams KH, Beller HR. 2016. Metatranscriptomic evidence of pervasive and diverse chemolithoautotrophy relevant to C, S, N and Fe cycling in a shallow alluvial aquifer. ISME J 10:2106–2117.

6. Trias R, Ménez B, le Campion P, Zivanovic Y, Lecourt L, Lecoeuvre A, Schmitt-Kopplin P, Uhl J, Gislason SR, Alfreðsson HA, Mesfin KG, Snæbjörnsdóttir SÓ, Aradóttir ES, Gunnarsson I, Matter JM, Stute M, Oelkers EH, Gérard E. 2017. High reactivity of deep biota under anthropogenic CO_2_ injection into basalt. Nat Commun 8:1–14.

7. McAllister SM, Polson SW, Butterfield DA, Glazer BT, Sylvan JB, Chan CS. 2020. Validating the Cyc2 neutrophilic iron oxidation pathway using meta-omics of Zetaproteobacteria iron mats at marine hydrothermal vents. mSystems 5:e00553–19.

8. McAllister SM, Moore RM, Gartman A, Luther GW III, Emerson D, Chan CS. 2019. The Fe(II)-oxidizing Zetaproteobacteria: historical, ecological and genomic perspectives. FEMS Microbiology Ecology 95:fiz015.

9. He S, Tominski C, Kappler A, Behrens S, Roden EE. 2016. Metagenomic analyses of the autotrophic Fe(II)-oxidizing, nitrate-reducing enrichment culture KS. Appl Environ Microbiol 82:2656–2668.

10. McAllister SM, Vandzura R, Keffer JL, Polson SW, Chan CS. 2020. Aerobic and anaerobic iron oxidizers together drive denitrification and carbon cycling at marine iron-rich hydrothermal vents. The ISME Journal https://doi.org/10.1038/s41396-020-00849-y.

11. Bird LJ, Bonnefoy V, Newman DK. 2011. Bioenergetic challenges of microbial iron metabolisms. Trends Microbiol 19:330–340.

12. Hedrich S, Schlömann M, Johnson DB. 2011. The iron-oxidizing proteobacteria. Microbiology (Reading) 157:1551–1564.

13. Ilbert M, Bonnefoy V. 2013. Insight into the evolution of the iron oxidation pathways. Biochim Biophys Acta 1827:161–175.

14. Moya-Beltrán A, Cárdenas JP, Covarrubias PC, Issotta F, Ossandon FJ, Grail BM, Holmes DS, Quatrini R, Johnson DB. 2014. Draft genome sequence of the nominated type strain of “*Ferrovum myxofaciens*,” an acidophilic, iron-oxidizing Betaproteobacterium. Genome Announc 2:1–2.

15. Quaiser A, Bodi X, Dufresne A, Naquin D, Francez A-J, Dheilly A, Coudouel S, Pedrot M, Vandenkoornhuyse P. 2014. Unraveling the stratification of an iron-oxidizing microbial mat by metatranscriptomics. PLOS ONE 9:1–9.

16. Fukushima J, Tojo F, Asano R, Kobayashi Y, Shimura Y, Okano K, Miyata N. 2015. Complete genome sequence of the unclassified iron-oxidizing, chemolithoautotrophic *Burkholderiales* bacterium GJ-E10, isolated from an acidic river. Genome Announc 3:1–2.

17. Fullerton H, Hager KW, McAllister SM, Moyer CL. 2017. Hidden diversity revealed by genome-resolved metagenomics of iron-oxidizing microbial mats from Lō’ihi Seamount, Hawai’i. The ISME Journal 11:1900–1914.

18. Ullrich SR, Poehlein A, Levicán G, Mühling M, Schlömann M. 2018. Iron targeted transcriptome study draws attention to novel redox protein candidates involved in ferrous iron oxidation in “*Ferrovum*” sp. JA12. Research in Microbiology 169:618–627.

19. Bethencourt L, Bochet O, Farasin J, Aquilina L, Borgne TL, Quaiser A, Biget M, Michon-Coudouel S, Labasque T, Dufresne A. 2020. Genome reconstruction reveals distinct assemblages of *Gallionellaceae* in surface and subsurface redox transition zones. FEMS Microbiology Ecology 96:fiaa036.

20. Cooper RE, Wegner C-E, McAllister SM, Shevchenko O, Chan CS, Küsel K. 2020. Draft genome sequence of *Sideroxydans* sp. strain CL21, an Fe(II)-oxidizing bacterium. Microbiol Resour Announc 9:e01444–19.

21. Emerson D, Field EK, Chertkov O, Davenport KW, Goodwin L, Munk C, Nolan M, Woyke T. 2013. Comparative genomics of freshwater Fe-oxidizing bacteria: implications for physiology, ecology, and systematics. Front Microbiol 4:254.

22. Kato S, Ohkuma M, Powell DH, Krepski ST, Oshima K, Hattori M, Shapiro N, Woyke T, Chan CS. 2015. Comparative genomic insights into ecophysiology of neutrophilic, microaerophilic iron oxidizing bacteria. Front Microbiol 6:1265.

23. Chiu BK, Kato S, McAllister SM, Field EK, Chan CS. 2017. Novel pelagic iron-oxidizing Zetaproteobacteria from the Chesapeake Bay oxic–anoxic transition zone. Front Microbiol 8:1280.

24. Crowe SA, Hahn AS, Morgan-Lang C, Thompson KJ, Simister RL, Llirós M, Hirst M, Hallam SJ. 2017. Draft genome sequence of the pelagic photoferrotroph *Chlorobium phaeoferrooxidans*. Genome Announc 5:e01584–16.

25. Mori JF, Scott JJ, Hager KW, Moyer CL, Küsel K, Emerson D. 2017. Physiological and ecological implications of an iron- or hydrogen-oxidizing member of the Zetaproteobacteria, Ghiorsea bivora, gen. nov., sp. nov. The ISME Journal 11:2624–2636.

26. Yarzábal A, Brasseur G, Ratouchniak J, Lund K, Lemesle-Meunier D, DeMoss JA, Bonnefoy V. 2002. The high-molecular-weight cytochrome c Cyc2 of *Acidithiobacillus ferrooxidans* is an outer membrane protein. J Bacteriol 184:313–317.

27. Castelle C, Guiral M, Malarte G, Ledgham F, Leroy G, Brugna M, Giudici-Orticoni M-T. 2008. A new iron-oxidizing/O_2_-reducing supercomplex spanning both inner and outer membranes, isolated from the extreme acidophile *Acidithiobacillus ferrooxidans*. J Biol Chem 283:25803–25811.

28. He S, Barco RA, Emerson D, Roden EE. 2017. Comparative genomic analysis of neutrophilic iron(II) oxidizer genomes for candidate genes in extracellular electron transfer. Front Microbiol 8:1584.

29. Akob DM, Hallenbeck M, Beulig F, Fabisch M, Küsel K, Keffer JL, Woyke T, Shapiro N, Lapidus A, Klenk H-P, Chan CS. 2020. Mixotrophic iron-oxidizing *Thiomonas* isolates from an acid mine drainage-affected creek. Appl Environ Microbiol 86:e01424–20.

30. Jiao Y, Newman DK. 2007. The *pio* operon is essential for phototrophic Fe(II) oxidation in Rhodopseudomonas palustris TIE-1. JB 189:1765-1773.

31. Liu J, Wang Z, Belchik SM, Edwards MJ, Liu C, Kennedy DW, Merkley ED, Lipton MS, Butt JN, Richardson DJ, Zachara JM, Fredrickson JK, Rosso KM, Shi L. 2012. Identification and characterization of MtoA: A decaheme c-type cytochrome of the neutrophilic Fe(II)-oxidizing bacterium *Sideroxydans lithotrophicus* ES-1. Front Microbiol 3:37.

32. Gupta D, Sutherland MC, Rengasamy K, Meacham JM, Kranz RG, Bose A. 2019. Photoferrotrophs produce a PioAB electron conduit for extracellular electron uptake. mBio 10.

33. Jeans C, Singer SW, Chan CS, Verberkmoes NC, Shah M, Hettich RL, Banfield JF, Thelen MP. 2008. Cytochrome 572 is a conspicuous membrane protein with iron oxidation activity purified directly from a natural acidophilic microbial community. ISME J 2:542–550.

34. Straub KL, Benz M, Schink B. 2001. Iron metabolism in anoxic environments at near neutral pH. FEMS Microbiology Ecology 34:181–186.

35. Johnson DB, Kanao T, Hedrich S. 2012. Redox transformations of iron at extremely low pH: Fundamental and applied aspects. Front Microbiol 3:96.

36. Majzlan J. 2013. Minerals and aqueous species of iron and manganese as reactants and products of microbial metal respiration., p. 1–28. In Microbial Metal Respiration. Springer Berlin Heidelberg, Berlin, Heidelberg.

37. Emerson D, Rentz JA, Lilburn TG, Davis RE, Aldrich H, Chan C, Moyer CL. 2007. A novel lineage of Proteobacteria involved in formation of marine Fe-oxidizing microbial mat communities. PLOS ONE 2:e667.

38. Singer E, Emerson D, Webb EA, Barco RA, Kuenen JG, Nelson WC, Chan CS, Comolli LR, Ferriera S, Johnson J, Heidelberg JF, Edwards KJ. 2011. *Mariprofundus ferrooxydans* PV-1 the first genome of a marine Fe(II) oxidizing Zetaproteobacterium. PLOS ONE 6:e25386.

39. Barco RA, Emerson D, Sylvan JB, Orcutt BN, Meyers MEJ, Ramírez GA, Zhong JD, Edwards KJ. 2015. New insight into microbial iron oxidation as revealed by the proteomic profile of an obligate iron-oxidizing chemolithoautotroph. Appl Environ Microbiol 81:5927–5937.

40. Almagro Armenteros JJ, Tsirigos KD, Sønderby CK, Petersen TN, Winther O, Brunak S, von Heijne G, Nielsen H. 2019. SignalP 5.0 improves signal peptide predictions using deep neural networks. Nature Biotechnology 37:420–423.

41. Scott RA, Mauk AG. 1996. Cytochrome C: a multidisciplinary approach. University Science Books, Sausalito, CA.

42. Jones DT. 1999. Protein secondary structure prediction based on position-specific scoring matrices. J Mol Biol 292:195–202.

43. Buchan DWA, Jones DT. 2019. The PSIPRED Protein Analysis Workbench: 20 years on. Nucleic Acids Research 47:W402–W407.

44. Zimmermann L, Stephens A, Nam S-Z, Rau D, Kübler J, Lozajic M, Gabler F, Söding J, Lupas AN, Alva V. 2018. A completely reimplemented MPI Bioinformatics Toolkit with a new HHpred server at its core. Journal of Molecular Biology 430:2237–2243.

45. Gabler F, Nam S-Z, Till S, Mirdita M, Steinegger M, Söding J, Lupas AN, Alva V. 2020. Protein sequence analysis using the MPI Bioinformatics Toolkit. Current Protocols in Bioinformatics 72:e108.

46. Moraes TF, Bains M, Hancock REW, Strynadka NCJ. 2007. An arginine ladder in OprP mediates phosphate-specific transfer across the outer membrane. Nat Struct Mol Biol 14:85–87.

47. Modi N, Ganguly S, Bárcena-Uribarri I, Benz R, van den Berg B, Kleinekathöfer U. 2015. Structure, dynamics, and substrate specificity of the OprO porin from *Pseudomonas aeruginosa*. Biophys J 109:1429–1438.

48. Söding J, Biegert A, Lupas AN. 2005. The HHpred interactive server for protein homology detection and structure prediction. Nucleic Acids Res 33:W244–248.

49. Zhang Y. 2008. I-TASSER server for protein 3D structure prediction. BMC Bioinformatics 9:40.

50. Kelley LA, Mezulis S, Yates CM, Wass MN, Sternberg MJE. 2015. The Phyre2 web portal for protein modeling, prediction and analysis. Nature Protocols 10:845–858.

51. Webb B, Sali A. 2016. Comparative protein structure modeling using MODELLER. Curr Protoc Bioinformatics 54:5.6.1–5.6.37.

52. Hancock RE. 1987. Role of porins in outer membrane permeability. J Bacteriol 169:929– 933.

53. Chan CS, Fakra SC, Emerson D, Fleming EJ, Edwards KJ. 2011. Lithotrophic iron-oxidizing bacteria produce organic stalks to control mineral growth: implications for biosignature formation. The ISME Journal 5:717–727.

54. Comolli LR, Luef B, Chan CS. 2011. High-resolution 2D and 3D cryo-TEM reveals structural adaptations of two stalk-forming bacteria to an Fe-oxidizing lifestyle. Environ Microbiol 13:2915–2929.

55. Hagan CL, Silhavy TJ, Kahne D. 2011. β-Barrel membrane protein assembly by the Bam complex. Annu Rev Biochem 80:189–210.

56. Noinaj N, Mayclin S, Stanley AM, Jao CC, Buchanan SK. 2016. From constructs to crystals - towards structure determination of β-barrel outer membrane proteins. J Vis Exp https://doi.org/10.3791/53245.

57. Londer YY. 2011. Expression of recombinant cytochromes c in E. coli, p. 123–150. In Evans, Jr., T, Xu, MQ (eds.), Heterologous gene expression in E. coli. Methods in Molecular Biology (Methods and Protocols). Humana Press.

58. Hancock RE, Carey AM. 1979. Outer membrane of *Pseudomonas aeruginosa*: heat- 2- mercaptoethanol-modifiable proteins. Journal of Bacteriology 140:902–910.

59. Hancock REW. 2021. Separation of outer and inner membranes. Hancock Laboratory Methods.

60. Gennis RB. 1987. The cytochromes of *Escherichia coli*. FEMS Microbiology Reviews 3:387–399.

61. Gustaffson JP. 2014. Visual MINTEQ. https://vminteq.lwr.kth.se/.

62. Massey V. 1991. Flavins and flavoproteins. Walter de Gruyter & Co., New York.

63. Efimov I, Parkin G, Millett ES, Glenday J, Chan CK, Weedon H, Randhawa H, Basran J, Raven EL. 2014. A simple method for the determination of reduction potentials in heme proteins. FEBS Letters 588:701–704.

64. Sutherland MC, Rankin JA, Kranz RG. 2016. Heme trafficking and modifications during System I cytochrome c biogenesis: Insights from heme redox potentials of Ccm proteins. Biochemistry 55:3150–3156.

65. White GF, Edwards MJ, Gomez-Perez L, Richardson DJ, Butt JN, Clarke TA. 2016. Chapter three - mechanisms of bacterial extracellular electron exchange, p. 87–138. In Poole, RK (ed.), Advances in Microbial Physiology. Academic Press.

66. Edwards MJ, White GF, Butt JN, Richardson DJ, Clarke TA. 2020. The crystal structure of a biological insulated transmembrane molecular wire. Cell 181:665–673.e10.

67. Shi L, Rosso KM, Zachara JM, Fredrickson JK. 2012. Mtr extracellular electron-transfer pathways in Fe(III)-reducing or Fe(II)-oxidizing bacteria: a genomic perspective. Biochemical Society Transactions 40:1261–1267.

68. Sanders C, Turkarslan S, Lee D-W, Daldal F. 2010. Cytochrome c biogenesis: the Ccm system. Trends in Microbiology 18:266–274.

69. Mayo SL, Ellis WR, Crutchley RJ, Gray HB. 1986. Long-range electron transfer in heme proteins. Science 233:948–952.

70. Nikaido H. 2003. Molecular basis of bacterial outer membrane permeability revisited. Microbiol Mol Biol Rev 67:593–656.

71. Beckwith CR, Edwards MJ, Lawes M, Shi L, Butt JN, Richardson DJ, Clarke TA. 2015. Characterization of MtoD from *Sideroxydans lithotrophicus*: a cytochrome c electron shuttle used in lithoautotrophic growth. Front Microbiol 6.

72. Abergel C, Nitschke W, Malarte G, Bruschi M, Claverie J-M, Giudici-Orticoni M-T. 2003. The structure of *Acidithiobacillus ferrooxidans* c4-cytochrome: A model for complex-induced electron transfer tuning. Structure 11:547–555.

73. Garber AI, Nealson KH, Okamoto A, McAllister SM, Chan CS, Barco RA, Merino N. 2020. FeGenie: A comprehensive tool for the identification of iron genes and iron gene neighborhoods in genome and metagenome assemblies. Front Microbiol 11:37.

74. Singer SW. 2012. Targeted isolation of proteins from natural microbial communities living in an extreme environment, p. In Navid, A (ed.), Microbial Systems Biology Methods and Protocols. Humana Press.

75. Jain A, Gralnick JA. 2021. Engineering lithoheterotrophy in an obligate chemolithoautotrophic Fe(II) oxidizing bacterium. Scientific Reports 11:2165.

76. Schmidt TG, Koepke J, Frank R, Skerra A. 1996. Molecular interaction between the Strep-tag affinity peptide and its cognate target, streptavidin. J Mol Biol 255:753–766.

77. Arslan E, Schulz H, Zufferey R, Künzler P, Thöny-Meyer L. 1998. Overproduction of the *Bradyrhizobium japonicum* c-type cytochrome subunits of the *cbb3* oxidase in *Escherichia coli*. Biochemical and Biophysical Research Communications 251:744–747.

78. Baron D, LaBelle E, Coursolle D, Gralnick JA, Bond DR. 2009. Electrochemical measurement of electron transfer kinetics by *Shewanella oneidensis* MR-1. Journal of Biological Chemistry 284:28865–28873.

79. Doss M, Philipp-Dormston WK. 1971. Porphyrin and heme biosynthesis from endogenous and exogenous δ-aminolevulinic acid in *Escherichia coli*, *Pseudomonas aeruginosa*, and *Achromobacter metalcaligenes*. Biological Chemistry 352:725–733.

80. Sudhamsu J, Kabir M, Airola MV, Patel BA, Yeh S-R, Rousseau DL, Crane BR. 2010. Co-expression of ferrochelatase allows for complete heme incorporation into recombinant proteins produced in *E. coli*. Protein Expr Purif 73:78–82.

81. Laemmli UK. 1970. Cleavage of structural proteins during the assembly of the head of bacteriophage T4. Nature 227:680–685.

82. Francis RT, Becker RR. 1984. Specific indication of hemoproteins in polyacrylamide gels using a double-staining process. Analytical Biochemistry 136:509–514.

83. Carlson HK, Clark IC, Blazewicz SJ, Iavarone AT, Coates JD. 2013. Fe(II) oxidation is an innate capability of nitrate-reducing bacteria that involves abiotic and biotic reactions. Journal of Bacteriology 195:3260–3268.

84. Finehout EJ, Lee KH. 2003. Comparison of automated in-gel digest methods for femtomole level samples. ELECTROPHORESIS 24:3508–3516.

85. Camacho C, Coulouris G, Avagyan V, Ma N, Papadopoulos J, Bealer K, Madden TL. 2009. BLAST+: architecture and applications. BMC Bioinformatics 10:421.

86. Li W, Godzik A. 2006. Cd-hit: a fast program for clustering and comparing large sets of protein or nucleotide sequences. Bioinformatics 22:1658–1659.

87. Edgar RC. 2004. MUSCLE: multiple sequence alignment with high accuracy and high throughput. Nucleic Acids Res 32:1792–1797.

88. Stamatakis A. 2014. RAxML version 8: a tool for phylogenetic analysis and post-analysis of large phylogenies. Bioinformatics 30:1312–1313.

89. Moore RM, Harrison AO, McAllister SM, Polson SW, Wommack KE. 2020. Iroki: automatic customization and visualization of phylogenetic trees. PeerJ 8:e8584.

90. Johnson LS, Eddy SR, Portugaly E. 2010. Hidden Markov model speed heuristic and iterative HMM search procedure. BMC Bioinformatics 11:431.

