## Supplemental Figures and Tables for "Iron oxidation by a fused cytochrome-porin common to diverse iron-oxidizing bacteria"

Running title: Neutrophilic iron-oxidizer Cyc2 is an iron oxidase

**Table of contents** 2
**Figure S1.** PSIPRED prediction of secondary structure 3
**Figure S2.** Constructs and expression of $Cyc2_{PV-1}$ 4
**Figure S3.** Uncropped gels 5
**Figure S4.** Four independent redox titration reactions 6
**Figure S5.** Three views of the modeled cytochrome domain of Cyc2PV-1 7
**Figure S6.** Full alignment of Cyc2 from neutrophilic and acidophilic FeOB 8
**Figure S7.** Histograms of pairwise amino acid identity 9
**Figure S8.** Comparison of motifs found in the conserved cytochrome domain 10
**Table S1.** Unique peptides detected by tandem MS/MS 11
**Table S2.** Relevant Fe(II) speciation 12

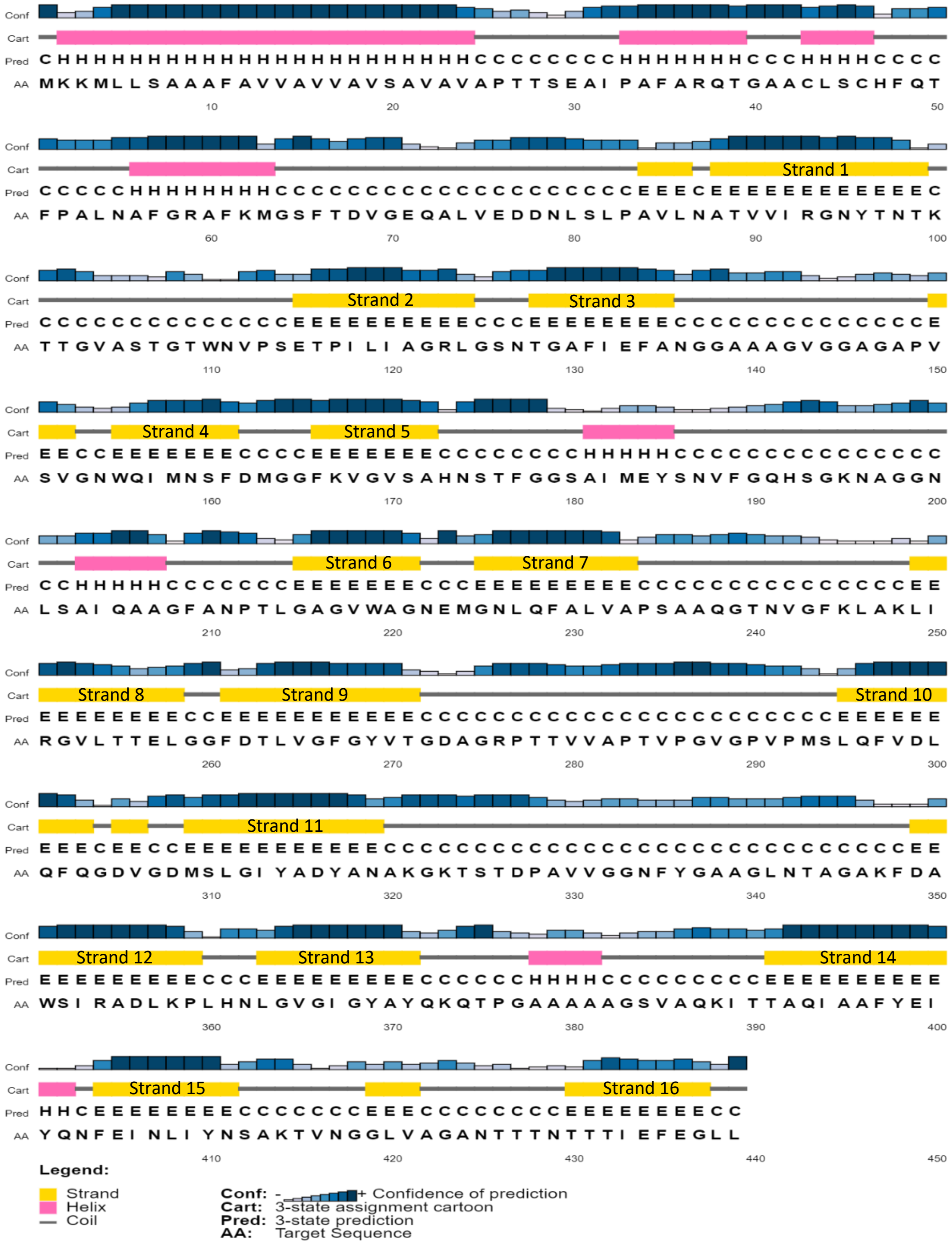


**Figure S1.** PSIPRED prediction of secondary structure in Cyc2$_{PV-1}$, the Cyc2 sequence from *M. ferrooxydans* PV-1 (D. W. A. Buchan, D. T. Jones, Nuc Acids Res, 2019, https://doi.org/10.1093/nar/gkz297).

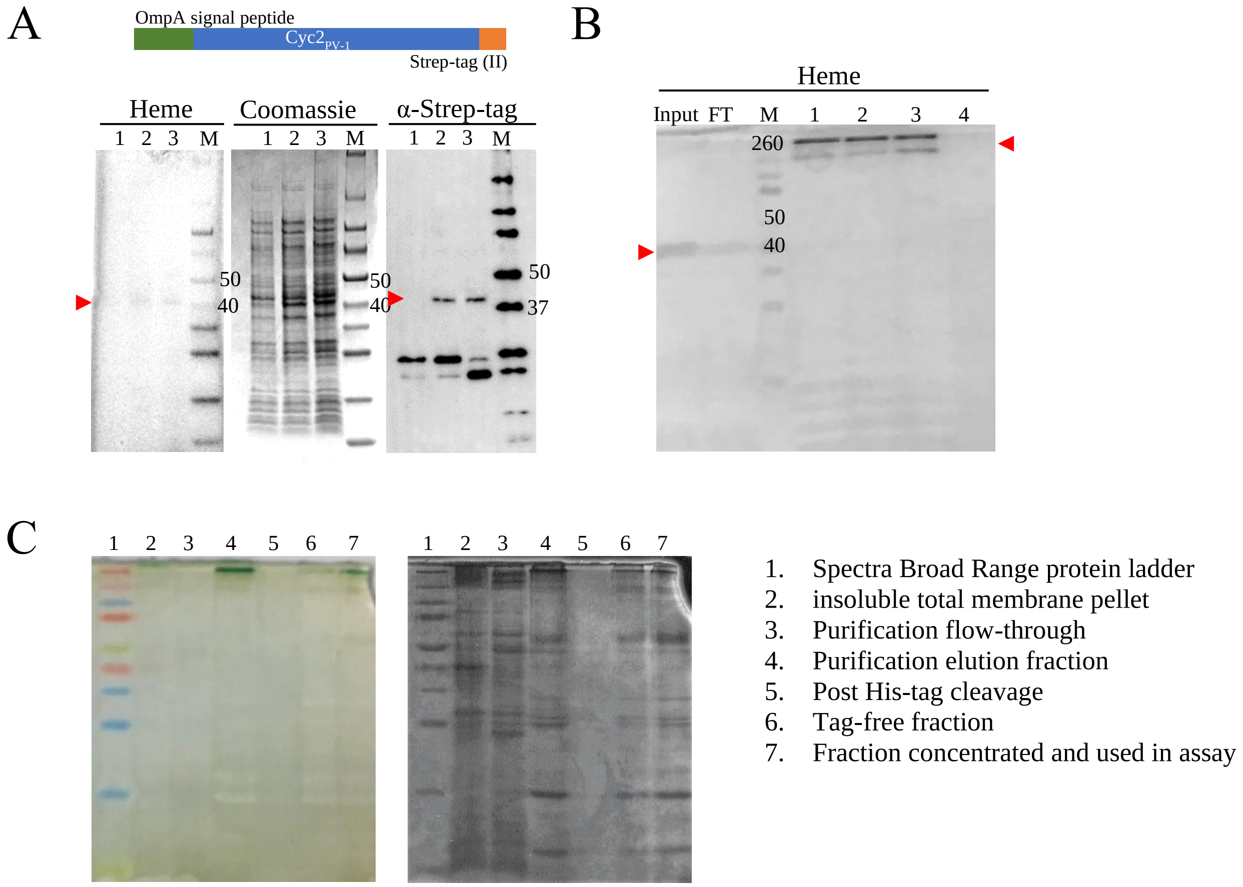


**Figure S2.** Constructs and expression of Cyc2$_{PV-1}$. Cyc2$_{PV-1}$ is marked with a red arrowhead, protein ladder is labeled with an M (Spectra Broad Range on heme and Coomassie, WesternC on α-*Strep*-tag Western blot), and relevant band sizes are labeled in kDa. (A) Schematic of gene construct for expression and representative stained SDS-PAGE gels showing Cyc2$_{PV-1}$ expression in *E. coli*: 1) uninduced, 2) induced, 3) lysed and induced. Smaller bands visible on *Strep*-tag Western blots are non-specific. (B) Heme-stained gel of fractions during His-tag purification. Cyc2$_{PV-1}$ migrates at its expected molecular weight in the diluted total membranes (input) and flow-though (FT). After elution, Cyc2$_{PV-1}$ migrates in a high-molecular weight complex (1 - 300 μM imidazole, 2 - after dialysis to remove imidazole). After TEV protease cleavage of the His-tag, Cyc2$_{PV-1}$ does not interact with the Ni-NTA column (3 - flow-through, 4 - imidazole elution). (C) Uncropped gel corresponding to Figure 2D. See lane labels to the right of image.

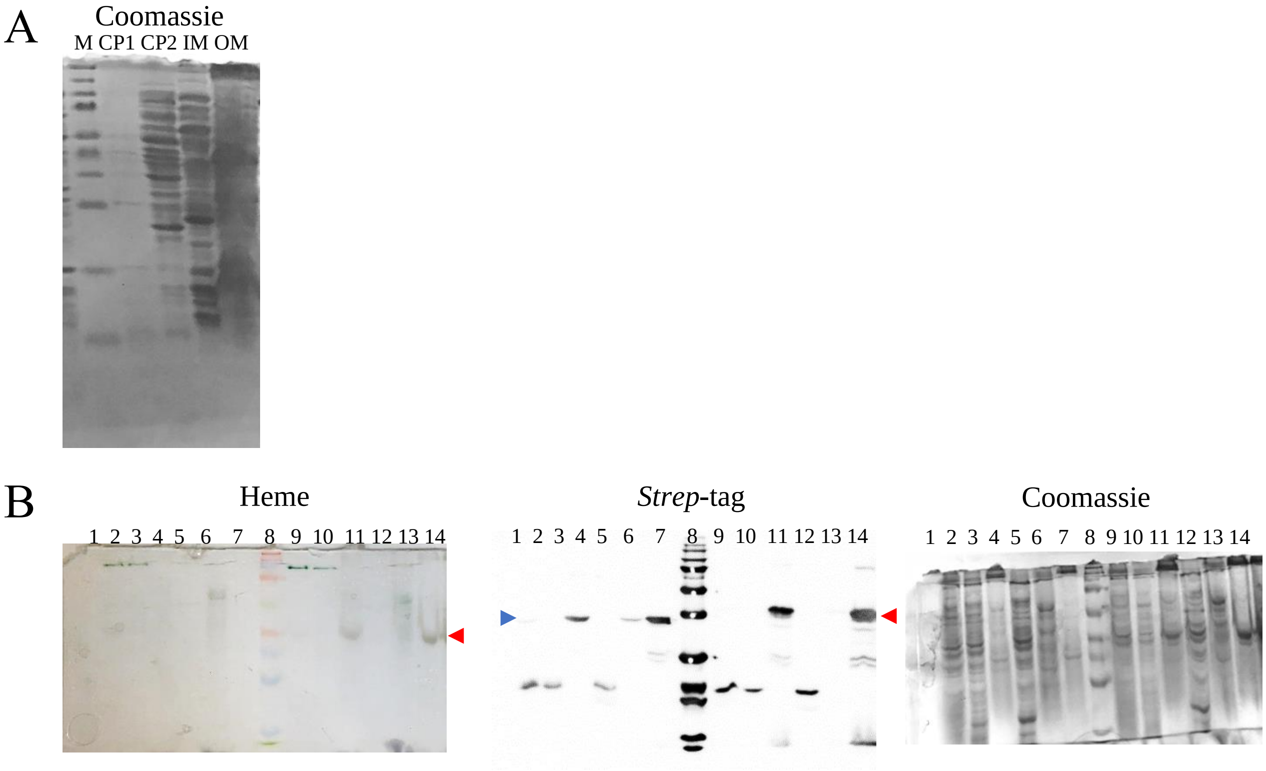


**Figure S3.** Uncropped gels. (A) Coomassie-stained gel corresponding to Fig. 3A. CP1, CP2 - cytoplasmic proteins, IM - inner membranes, OM - outer membranes, M - Spectra Broad Range protein ladder with relevant bands in kDa. (B) Gels corresponding to samples in Fig. 4. Red arrow indicates full-length $Cyc2_{PV-1}$ and blue arrow indicates porin-only. Lanes: 1-empty vector, 2-porin lysed supernatant, 3-porin ultracentrifuged supernatant, 4-porin total membranes, 5-porin cytoplasmic proteins, 6-porin inner membranes, 7-porin outer membranes, 8-Spectra Broad Range or WesternC protein ladder, 9- $Cyc2_{PV-1}$ lysed supernatant, 10- $Cyc2_{PV-1}$ ultracentrifuged supernatant, 11- $Cyc2_{PV-1}$ total membranes, 12- $Cyc2_{PV-1}$ cytoplasmic proteins, 13- $Cyc2_{PV-1}$ inner membranes, 14- $Cyc2_{PV-1}$ outer membranes.

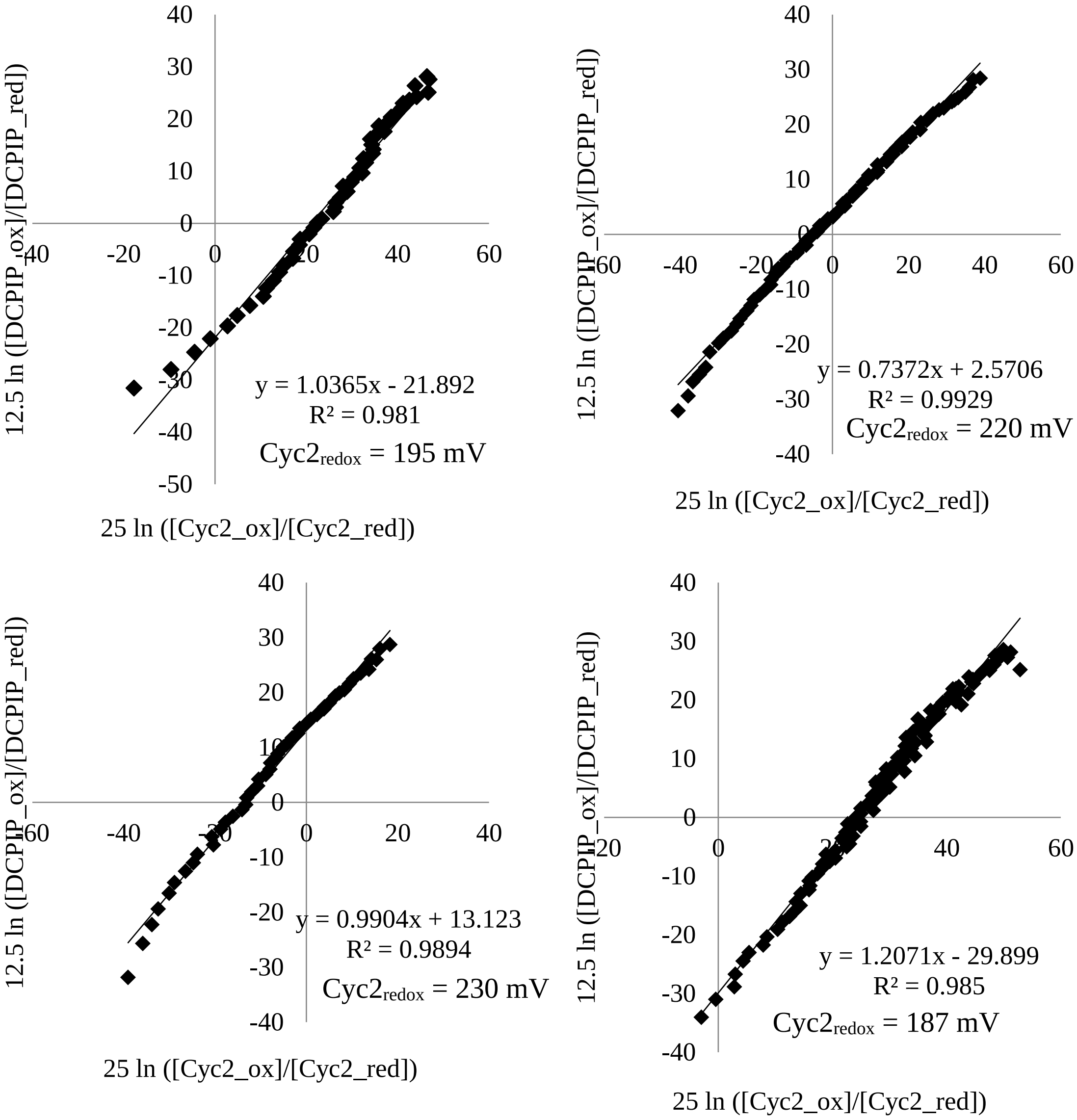


**Figure S4.** Four independent redox titration reactions with $Cyc2_{PV-1}$ and DCPIP.

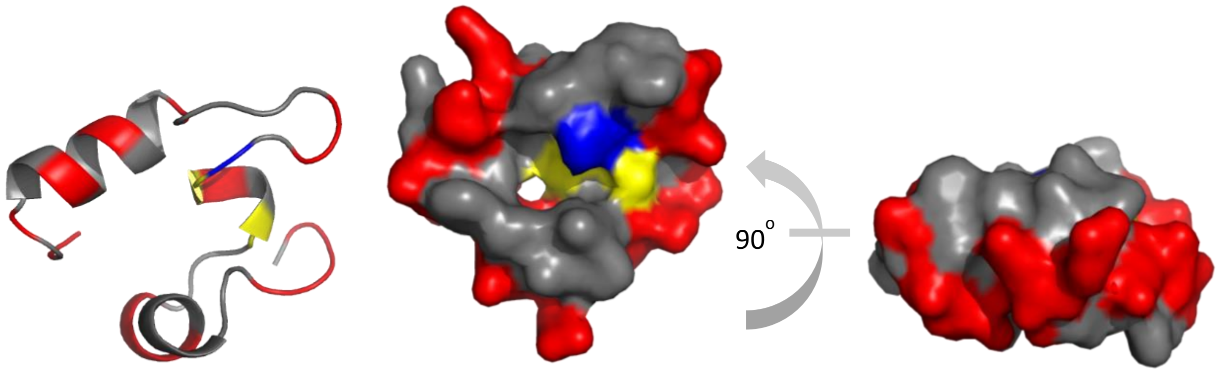


**Figure S5.** Three views of the modeled cytochrome domain of $Cyc2_{PV-1}$. The view on the right is rotated 90 $^{o}$ away from the viewer compared to the view in the center. Hydrophobic residues are gray and polar residues are red. Heme (not pictured) is covalently attached to cysteine residues (yellow) and coordinated by histidine (blue). Model generated using MODELLER (B. Webb, A. Sali, Curr Prot Bioinf, 54: 5.6.1-5.6.37, 2016, https://doi: 10.1002/cpbi.3).

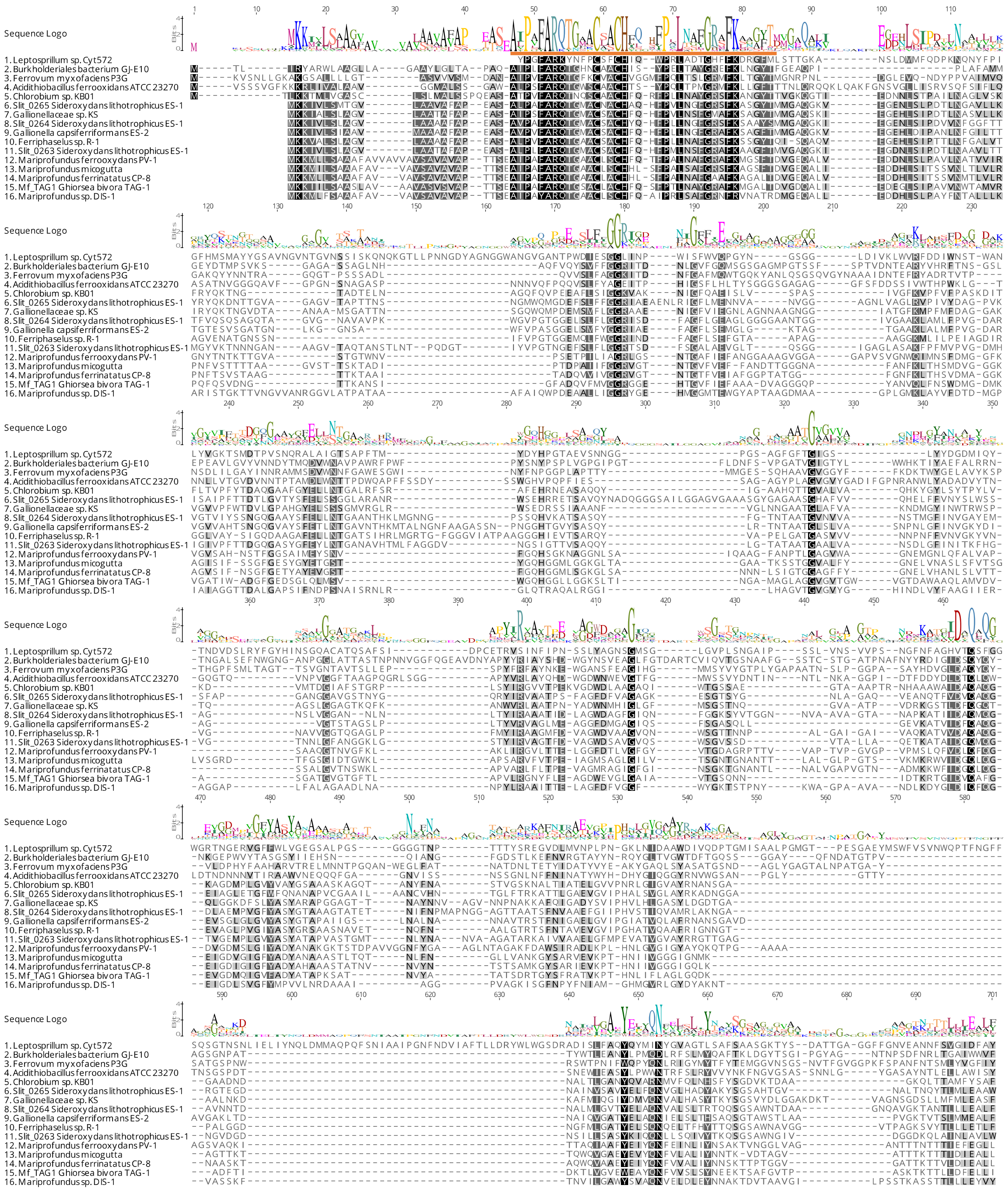


**Figure S6.** Full alignment of Cyc2 from representative neutrophilic and acidophilic FeOB. Orange line indicates the conserved cytochrome region.

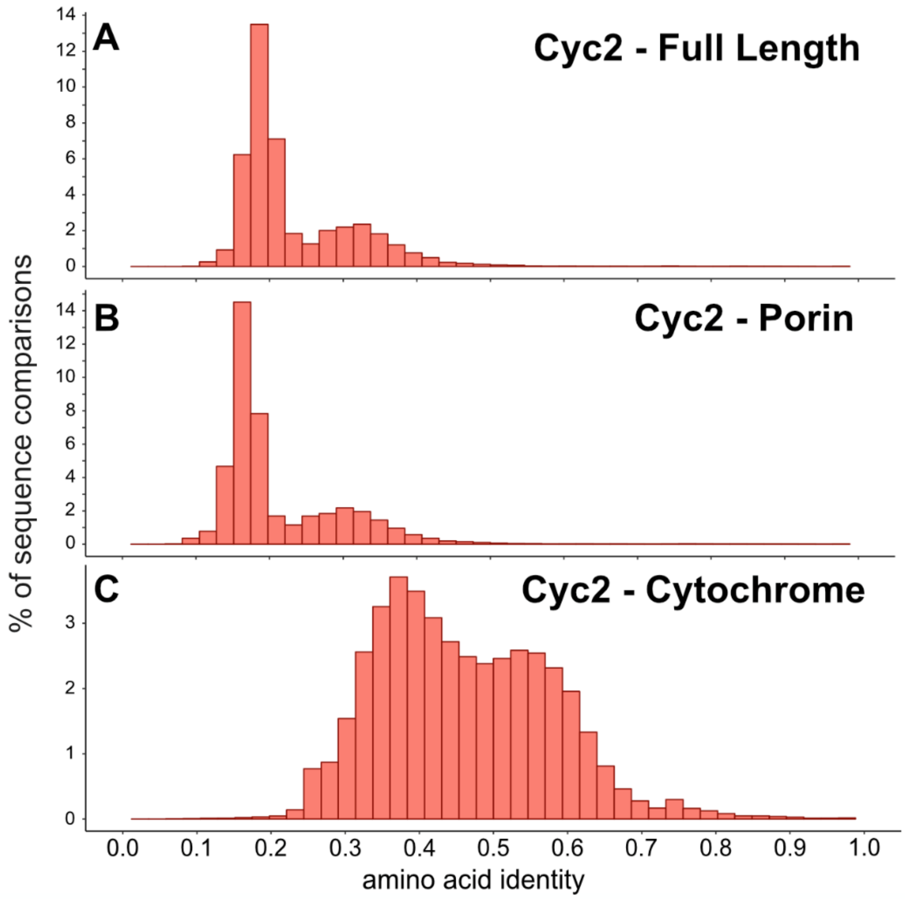


| | | Cluster 1 | | | | | | | | | Cluster 2 | | Cluster 3 | | | |
|---|---|---|---|---|---|---|---|---|---|---|---|---|---|---|---|---|
| | | Zetaproteobacteria | | | | Gallionellaceae | | | | Chlorob. | **Lepto** | Tenderia | Ferrovum | Dechlor. | Burkh. | **Acidith.** |
| Cluster | Taxa | TAG-1 | DIS-1 | PV-1 | CP-8 | KS | ES-1 | ES-2 | OYT1 | luteo. | **ferrodz.** | electro. | myxo. | RCB | GJ-E10 | **ferrox.** |
| 1 | Ghiorsea bivora TAG-1 | | | | | | | | | | | | | | | |
| | Mariprofundus sp. DIS-1 | 29.9% | | | | | | | | | | | | | | |
| | M. ferrooxydans PV-1 | 38.9% | 33.5% | | | | | | | | | | | | | |
| | M. ferrinatatus CP-8 | 47.8% | 33.1% | 45.7% | | | | | | | | | | | | |
| | Gallionellaceae sp. KS | 28.2% | 26.9% | 30.1% | 28.3% | | | | | | | | | | | |
| | Sideroxydans lithotrophicus ES-1 | 29.2% | 27.3% | 29.1% | 29.1% | 39.6% | | | | | | | | | | |
| | Gallionella capsiferriformans ES-2 | 30.4% | 26.4% | 29.5% | 30.6% | 36.8% | 53.4% | | | | | | | | | |
| | Ferriphaselus amnicola OYT1 | 29.9% | 28.8% | 29.6% | 28.1% | 34.2% | 51.1% | 52.2% | | | | | | | | |
| | Chlorobium luteolum | 25.1% | 26.2% | 28.3% | 26.3% | 35.2% | 33.5% | 32.7% | 33.5% | | | | | | | |
| 2 | **Leptospirillum ferrodiazotrophum** | **18.8%** | **17.5%** | **16.4%** | **15.2%** | **19.4%** | **19.8%** | **18.1%** | **16.5%** | **17.9%** | | | | | | |
| | Tenderia electrophaga | 19.0% | 17.5% | 21.5% | 18.5% | 19.7% | 19.2% | 17.6% | 17.8% | 20.4% | **17.5%** | | | | | |
| 3 | Ferrovum myxofaciens P3G | 19.9% | 19.4% | 22.0% | 19.5% | 22.1% | 22.6% | 20.1% | 19.6% | 19.6% | **19.6%** | 18.9% | | | | |
| | Dechloromonas aromatica RCB | 21.6% | 17.2% | 19.7% | 19.3% | 18.6% | 19.5% | 19.5% | 19.1% | 18.2% | **18.2%** | 17.8% | 26.8% | | | |
| | Burkholderiales bacterium GJ-E10 | 21.1% | 19.6% | 19.8% | 19.6% | 20.0% | 22.1% | 22.7% | 21.7% | 19.7% | **17.4%** | 17.8% | 30.5% | 27.3% | | |
| | **Acidithiobacillus ferrooxidans ATCC2327** | **18.0%** | **21.5%** | **19.6%** | **18.9%** | **22.0%** | **23.0%** | **21.3%** | **23.1%** | **20.0%** | **18.6%** | **19.3%** | **28.1%** | **26.7%** | **28.2%** | |
| | Thiomonas sp. FB-6 | 22.0% | 20.9% | 21.8% | 20.6% | 22.7% | 19.4% | 20.4% | 22.9% | 21.4% | **17.4%** | 19.1% | 29.2% | 35.2% | 30.5% | **29.6%** |

**Figure S7.** Histograms of pairwise amino acid identity of the (A) full length Cyc2 sequences, (B) porin portion, and (C) cytochrome portion (n=156). The cytochrome portion is more highly conserved than the porin. (D) Amino acid identities (AAI) of full length Cyc2 sequences from FeOB and *Tenderia electrophaga*. AAI to biochemically characterized Cyc2 are shown in bold. Note that organisms from "Cluster 1" e.g. neutrophilic FeOB Zetaproteobacteria, Gallionellaceae, and *Chlorobi* are most similar to one another.

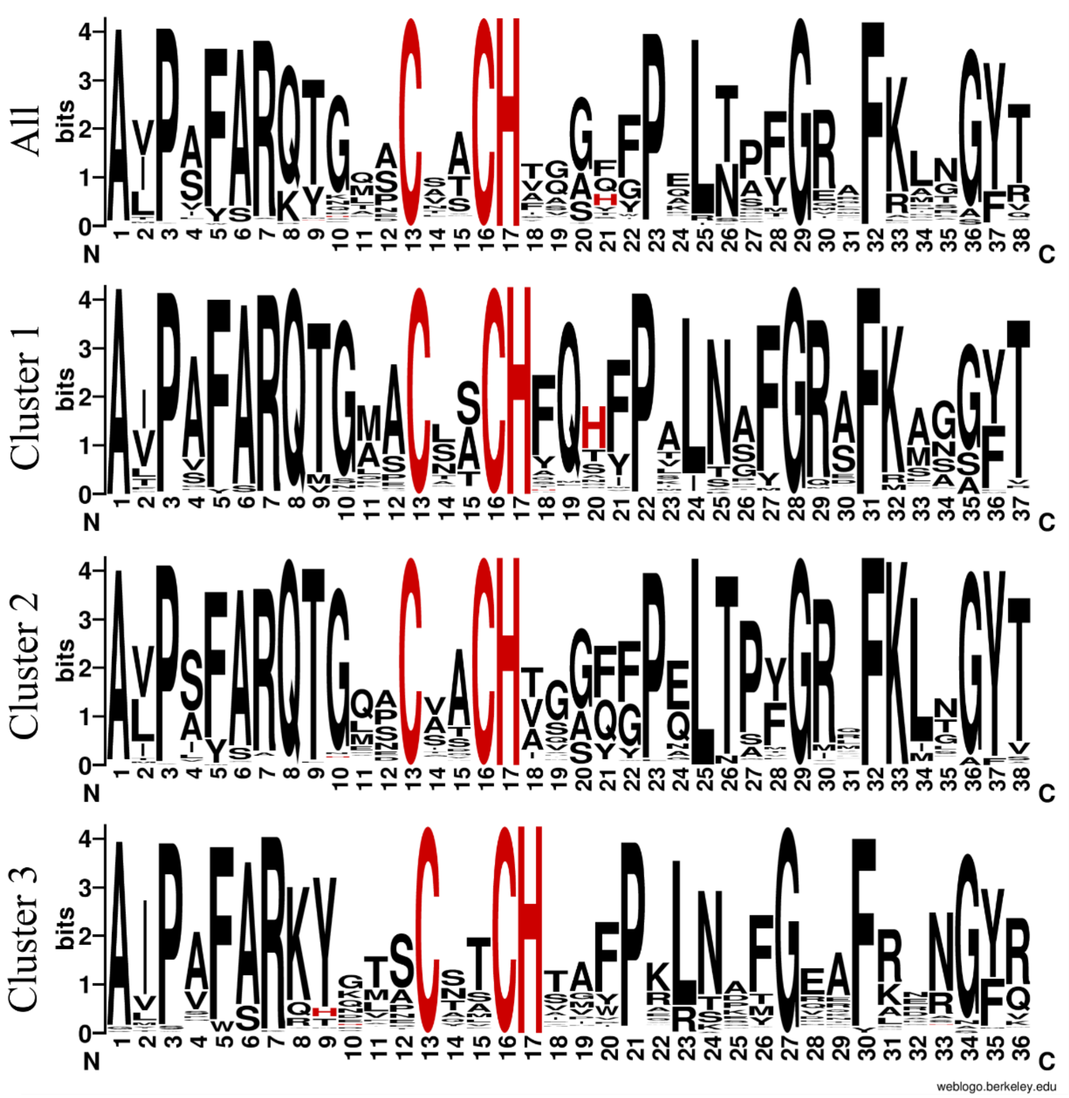


**Figure S8.** Comparison of motifs found in the conserved cytochrome domain of Cyc2. The sequence logo labeled "All" is built from 1593 homologs. Each of the Cluster logos are built from all sequences in each cluster (334 in Cluster 1, 858 in Cluster 2, and 401 in Cluster 3).

**Table S1.** Unique peptides detected by tandem MS/MS that matched to Cyc2$_{PV-1}$ with 99% confidence

| **Sequence** | **ΔMass** | **Obs. MW** | **Theor. MW** | **Modi-fications** |
|---|---|---|---|---|
| ADLKPLHNLGVGIGYAYQK | 3.24E-03 | 2056.114 | 2056.111 | |
| ADLKPLHNLGVGIGYAYQKQTPGAAAAAGSVAQK | -2.54E-03 | 3364.787 | 3364.789 | |
| FDAWSIR | 2.21E-03 | 893.4417 | 893.4396 | |
| GKTSTDPAVVGGNFYGAAGLNTAGAK | 5.05E-03 | 2423.213 | 2423.208 | |
| GVLTTELGGFDTLVGFGYVTGDAGR | -0.00189 | 2501.242 | 2501.244 | |
| ITTAQIAAFYEIYQNFEINLIYNSAK | 4.00E-03 | 3037.547 | 3037.544 | |
| MGSFTDVGEQALVEDDNLSLPAVLNATVVIR | -1.31E-03 | 3272.657 | 3272.66 | |
| PTTVVAPTVPGVGPVPM | -5.37E-04 | 1617.88 | 1617.88 | |
| QTGAACLSCHFQTFPALNAFGR | -1.99E-04 | 2453.137 | 2453.137 | *a* |
| QTPGAAAAAGSVAQK | 0.000334 | 1309.663 | 1309.663 | *b* |
| QTPGAAAAAGSVAQKITTAQIAAFYEIYQNFEINLIYNSAK | -3.02E-03 | 4346.22 | 4346.222 | |
| TSTDPAVVGGNFYGAAGLNTAGAK | -0.00012 | 2238.092 | 2238.092 | |
| TTGVASTGTWNVPSETPILIAGR | -0.00015 | 2331.207 | 2331.207 | *c* |
| TVNGGLVAGANTTTNTTTIEFEGLLWSHPQFEK | -0.00022 | 3532.748 | 3532.747 | |
| VGVSAHNSTFGGSAIMEYSNVFGQHSGK | -0.00157 | 2867.328 | 2867.33 | |

[a]Carbamidomethyl at C6 and C9

[b]Gln->pyro-Glu at N-term

[c]Trp->Kynurenin at W10

**Table S2.** Relevant Fe(II) and citrate speciation in the iron oxidase assay buffer[a] from Visual MINTEQ calculation (https://vminteq.lwr.kth.se/).

| Component | % of total concentration | Species name |
|---|---|---|
| Citrate$^{3-}$ | 9.559 | Citrate$^{3-}$ |
| | 0.505 | FeH-citrate (aq.) |
| | 10.261 | Na-citrate$^{2-}$ |
| | 2.539 | H-citrate$^{2-}$ |
| | 0.029 | H2-citrate$^{-}$ |
| | 77.107 | Fe-citrate$^{-}$ |
| $Fe^{2+}$ | 21.282 | $Fe^{2+}$ |
| | 0.505 | FeH-citrate (aq.) |
| | 10.99 | $FeCl^{+}$ |
| | 77.107 | Fe-citrate$^{-}$ |
| $Na^{+}$ | 93.288 | $Na^{+}$ |
| | 0.068 | Na-citrate$^{2-}$ |
| | 6.644 | NaCl (aq.) |
| $Cl^{-}$ | 93.393 | $Cl^{-}$ |
| | 6.6 | NaCl (aq.) |
| MES$^{-}$ | 60.951 | MES$^{-}$ |
| | 39.049 | H-MES (aq.) |

[a]Sucrose and DDM were in assay buffer but not available as part of the calculation
